# THE ASSOCIATION OF CD47 WITH INTEGRIN Mac-1 REGULATES MACROPHAGE RESPONSES BY STABILIZING THE EXTENDED INTEGRIN CONFORMATION

**DOI:** 10.1101/2022.09.30.510402

**Authors:** Nataly P. Podolnikova, Shundene Key, Xu Wang, Tatiana P. Ugarova

## Abstract

CD47 is a ubiquitously expressed cell surface integrin-associated protein. Recently, we have demonstrated that integrin Mac-1 (α_M_β_2_, CD11b/CD18, CR3), the major adhesion receptor on the surface of myeloid cells, can be coprecipitated with CD47. However, the molecular basis for the CD47-Mac-1 interaction and its functional consequences remain unclear. Here, we demonstrated that CD47 regulates macrophage functions directly interacting with Mac-1. In particular, adhesion, spreading, migration, phagocytosis, and fusion of CD47-deficient macrophages were significantly decreased. The functional link between CD47 and Mac-1 was validated by co-immunoprecipitation analysis using various Mac-1-expressing cells. In HEK293 cells expressing individual α_M_ and β_2_ integrin subunits, CD47 has been found to bind both subunits. Interestingly, the amount of CD47 recovered with the free β_2_ subunit was higher than in the complex with the whole integrin. Furthermore, activating Mac-1-expressing HEK293 cells with PMA, Mn^2+^, and activating antibody increased CD47 in complex with Mac-1, suggesting greater stability of the complex with integrin in the extended conformation. Notably, on the surface of cells lacking CD47, fewer Mac-1 molecules could convert into an extended conformation in response to activation. The binding site in CD47 for Mac-1 was identified in its constituent IgV domain. The complementary binding sites for CD47 in Mac-1 were localized in integrin epidermal growth factor-like domains 3 and 4 of the β_2_ and calf-1 and calf-2 domains of the α subunits. These results indicate that Mac-1 forms a lateral complex with CD47, which regulates essential macrophage functions by stabilizing the extended integrin conformation.

## INTRODUCTION

CD47 is a ubiquitously expressed transmembrane protein implicated as an essential regulator of integrin function and a marker of “self” in normal physiological processes (1–3). It is also involved in many pathophysiological processes, such as inflammation, tumorigenesis, and others (4). CD47 is a member of the immunoglobulin (Ig) superfamily with a single extracellular IgV-like domain and five-pass transmembrane segments. CD47 has been initially identified as a ~50 kDa protein copurified with integrin α_v_β_3_ from neutrophils, platelets, and placenta and named integrin-associated protein (IAP) (5–7). Early *in vitro* studies showed that CD47 is required for numerous α_v_β_3_-dependent functions of neutrophils. In particular, function-blocking anti-CD47 antibodies reduced binding of vitronectin-coated beads, Fc-receptor-mediated phagocytosis, chemotaxis, migration through endothelial and epithelial barriers, increase in the concentration of intracellular Ca^2+^, and oxidative burst (6–12). Subsequent studies in CD47^-/-^ mice using several animal models of inflammation demonstrated that CD47 was involved in transendothelial migration of leukocytes (13,14). In addition to α_v_β_3_, CD47 has been shown to associate with integrins α_2_β_1_ and α_IIIb_β_3_ on platelets (15,16), α_2_β_1_ on smooth muscle cells (17), α_4_β_1_ on sickle erythrocytes (18), and α_5_β_1_ in chondrocytes (19). CD47 has also been found to regulate adhesive functions of α_L_β_2_ (20), a member of the β_2_ subfamily of integrins expressed on leukocytes, and the interaction between α_L_β_2_ and CD47 on the surface of cultured T-cells was detected by fluorescence lifetime imaging microscopy (20). Studies of the molecular requirements for the association of CD47 with integrin α_v_β_3_ have implicated the IgV domain of CD47 since GPI-linked IgV expressed on the surface of CD47-deficient ovarian cancer cells was capable of restoring the α_v_β_3_-mediated vitronectin binding (12). Furthermore, chimeric molecules containing only IgV and the first transmembrane segment of CD47 were sufficient to restore the arrest of CD47-negative Jurkat T-cells on VCAM-1 (21).

Although the exact mechanism by which CD47 regulates integrin function remains elusive, the association of CD47 with integrins that belong to different subfamilies and are expressed in various cells suggests that this interaction is required for a common process. We postulated that CD47 associates with and modulates functions not only of α_L_β_2_ but other leukocyte β_2_ integrins, in particular, integrin Mac-1 (α_M_β_2_, CD11b/CD18, CR3). Indeed, we have recently shown that Mac-1 can be coprecipitated with CD47 from Mac-1-expressing HEK293 cells and RAW264.7 murine macrophages (22). Among the β_2_ integrins, Mac-1 is the most abundant and versatile receptor on the surface of myeloid leukocytes. It mediates numerous adhesive reactions of neutrophils and monocyte/macrophages during the inflammatory response (23,24). In particular, it contributes to the firm adhesion of neutrophils to endothelial cells, promotes their diapedesis, and participates in migration of neutrophils to sites of inflammation. Ligand engagement by Mac-1 initiates various other leukocyte responses, including phagocytosis, respiratory burst, homotypic aggregation, and degranulation.

Like other β_2_ integrin subfamily members, Mac-1 is a heterodimeric receptor composed of the α subunit (CD11b) and the common β_2_ subunit (CD18). The α_M_ subunit contains an inserted I-domain, a region of ~200 amino acid residues, a characteristic feature of β_2_ integrins responsible for the binding of multiple ligands (25). Like other integrins, activation of Mac-1 during the immune-inflammatory response is accompanied by conformational changes resulting in the conversion of integrin from the bent to the extended conformation, which is detected by conformation-sensitive antibodies (26). Despite the critical role of Mac-1 in leukocyte biology, the functional significance of its association with CD47 and the molecular basis for the CD47-Mac-1 interaction remains unexplored.

In the present study, we investigated the effect of CD47 deficiency on Mac-1-mediated macrophage responses and determined the mechanism by which CD47 modulates integrin’s function. This study also aimed to identify the region in CD47 responsible for Mac-1 binding and the complementary site(s) for CD47 in Mac-1. We demonstrated that the Mac-1-CD47 complex is required for numerous Mac-1-dependent macrophage responses, including adhesion, migration, spreading, phagocytosis and fusion. The conversion of Mac-1 from the bent to extended conformation upon treatment of cells with activating stimuli increased the amount of CD47 in the Mac-1-CD47 complexes, suggesting that CD47 has an increased affinity for integrin in the extended conformation. The studies with CD47-deficient cells showed that CD47 is necessary to facilitate the extension of Mac-1. The IgV domain of CD47 and the integrin epidermal growth factor-like domains 3 and 4 (EGF-3 and EGF-4) of the β_2_ subunit and calf-1 and calf-2 domains of the α_M_ subunit have been identified as complementary binding sites. These findings are significant as they provide evidence for the direct interaction of CD47 with Mac-1 and present a potential for selective modulation of integrin function.

## RESULTS

### CD47 regulates Mac-1-dependent macrophage responses

To explore how CD47 might affect the Mac-1 function, we investigated the effect of CD47 deficiency on various macrophage responses, including adhesion, spreading, migration, phagocytosis, and IL-4-mediated macrophage fusion. In this set of experiments, we examined the reactions known to depend specifically on Mac-1 but not other integrins. Figures 1A and B show that adhesion of CD47-deficient macrophages to fibrinogen and ICAM-1, biologically relevant Mac-1 ligands (27,28), was significantly reduced compared to wild-type (WT) macrophages. At 2.5-5 μg/ml, the concentrations of fibrinogen that mediate the maximal Mac-1-mediated adhesion of many cell types (29,30), adhesion of CD47-deficient macrophages was ~2-fold less than that of WT cells. Similarly, adhesion of CD47-deficient cells to ICAM-1 was reduced by ~2.4-fold. The effect of CD47 deficiency was not due to the difference in the surface expression level of Mac-1, which was similar in WT and CD47-deficient macrophages (Fig. S1). In agreement with the cell adhesion results, the spreading of CD47-deficient macrophages on fibrinogen and ICAM-1 was significantly lower than WT counterparts (Fig. 1, C-E). Furthermore, unlike WT macrophages, CD47-deficient cells lacked podosomes, the actin-based structures containing integrins, talin, vinculin, and other proteins (Fig. 1C, arrowheads).

**Figure 1.**
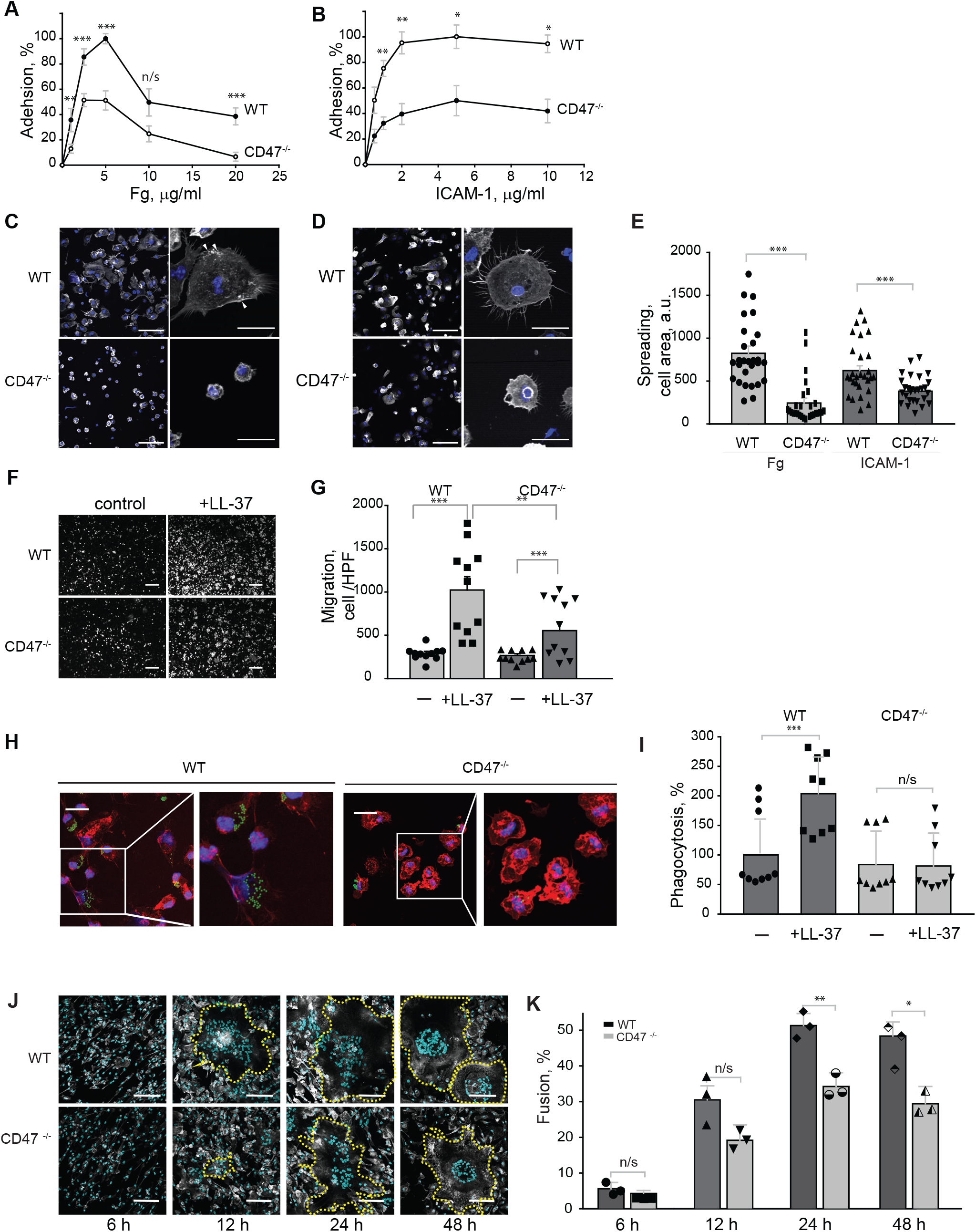
Requirement of CD47 in mediating Mac-1-dependent macrophage responses. Inflammatory macrophages were isolated from the peritoneum of wild-type and CD47^-/-^ mice on day 3 after TG injection and purified as described in the Experimental Procedures. **(A, B)** Effect of CD47 deficiency on macrophage adhesion. Aliquots of calcein-labeled macrophages were added to microtiter wells coated with different concentrations of fibrinogen (A) and ICAM-1 (B). After 30 min at 37 °C, nonadherent cells were removed by washing, and the fluorescence of adherent cells was measured. Data are expressed as a percentage of maximal adhesion determined at 5 μg/ml of each ligand and are means ± SE from 3 experiments with triplicate measurements. **(C-E)** Effect of CD47 deficiency on macrophage spreading. Macrophages were added to glass coverslips coated with 2.5 μg/ml fibrinogen (C) or 2 μg/ml ICAM-1 (D) and allowed to adhere and spread for 1 h at 37 °C. Samples were fixed and incubated with Alexa Fluor 488-conjugated phalloidin to detect F-actin (gray) and DAPI to stain nuclei (blue). The scale bars are 20 μm. The cell area was determined from confocal images of 30 cells adherent and spread on each ligand using ImageJ software and expressed in arbitrary units (a.u.) (E). Data shown are means ± SE from three individual experiments. Arrowheads indicate the podosomes. **(F, G)** Effect of CD47 deficiency on macrophage migration. Macrophages isolated from the inflamed peritoneum of WT and CD47^-/-^ mice were added to the upper chamber of the Transwell system, and the cathelicidin peptide LL-37 (5 μg/ml) was added to the bottom chamber. Macrophages were allowed to migrate for 90 min at 37 °C. Representative images of fluorescent cells that migrated through the filter of the Transwell system and remained attached to the underside of the filter are shown in (F). The number of cells per high power field (600 mm^2^) was counted (G). Data are presented as migrated cells per field ± SE for five random 20x fields per chamber from three individual experiments. **(H, I)** The CD47 requirement for Mac-1-dependent phagocytosis of opsonized beads. Fluorescent latex beads were preincubated with LL-37 (40 μg/ml) for 30 min at 37 °C, and soluble LL-37 was removed from beads by high-speed centrifugation. LL-37-coated beads were incubated with adherent mouse peritoneal macrophages isolated from WT and CD47^-/-^ mice for 30 min at 37 °C. Non-phagocytosed beads were removed, and phagocytosis was determined. Representative images of WT and CD47-deficient macrophages after uptake of fluorescent beads are shown in (H). Phagocytosis was quantified from ten fields of fluorescent images (I). The scale bars are 50 μM. Data shown are means ± SE from three experiments. **(J, K)** Effect of CD47 deficiency on macrophage fusion. Inflammatory macrophages isolated from WT and CD47^-/-^ mice were seeded on coverslips, and fusion was induced with IL-4. After different times, cells were fixed and stained with Alexa Fluor 647-conjugated phalloidin and DAPI (J). Fusion indices were calculated from confocal images as described in Experimental Procedures (K). Ten images were analyzed per time point. Data shown are means ± SE from three individual experiments. *p< .05, ** p< .01, ***p< .001.

It is known that *in vitro* Mac-1 promotes migration of macrophages toward its ligands (31–33). To determine whether CD47 deficiency affects the ability of Mac-1 to participate in macrophage migration, we used the cathelicidin peptide LL-37, a Mac-1 ligand that mediates migration of WT but not Mac-1-deficient macrophages (32). As shown in Fig. 1, F and G, migration of CD47-deficient macrophages to LL-37 in a Transwell system was significantly decreased (by 1.8-fold) compared to WT cells.

Another important function of Mac-1 is phagocytosis (23,24). Previous studies demonstrated that Mac-1 is involved in mediating phagocytosis of latex beads opsonized with various Mac-1 ligands, including LL-37 (32). The effect of CD47 deficiency on the ability of macrophages to phagocytose opsonized particles was examined by incubating adherent macrophages isolated from WT and CD47^-/-^ mice with LL-37-coated beads and determining the phagocytosis index. The opsonin-mediated phagocytosis by CD47-deficient macrophages was strongly impaired (Fig. 1, H and I). In particular, while phagocytosis of LL-37-coated beads by WT macrophages increased ~2-fold compared to uncoated beads, CD47-deficient macrophages ingested beads to the same extent as uncoated beads, suggesting that Mac-1 lost its ability to participate in opsonin-dependent phagocytosis.

As an adhesion receptor, Mac-1 is involved in macrophage fusion induced by IL-4 *in vitro* (34,35). The role of CD47 in this process was examined by plating inflammatory macrophages isolated from WT and CD47^-/-^ mice on the fusion-promoting substrate and determining the kinetics of fusion. Figure 1 (J and K) shows that CD47 was required for macrophage fusion inasmuch as the formation of multinucleated giant cells from CD47-deficient macrophages was significantly reduced at all tested times.

### Mac-1 and CD47 associate on the surface of Mac-1-expressing cells

We previously reported that CD47 could be immunoprecipitated with Mac-1 from suspended Mac-1-expressing HEK293 cells that endogenously express CD47 and murine RAW264.7 macrophages (22). We performed additional immunoprecipitation experiments to corroborate this finding, extend it to naturally occurring macrophages, and examine whether Mac-1 and CD47 form a complex on the surface of adherent cells. In agreement with previous data in RAW265.7 macrophages, mAbs M1/70 against the mouse α_M_ integrin subunit immunoprecipitated Mac-1 and CD47 from isolated peritoneal mouse macrophages and a mouse macrophage cell line IC-21 (Fig. 2, A, B). Control immunoprecipitations performed with isotype-specific IgG did not pull down these proteins. Furthermore, consistent with previous data, CD47 was found in complex with integrin in Mac-1-HEK293 cells (Fig. 2C). As determined by ratios of CD47 to the α_M_ and β_2_ integrin subunits present in the immunoprecipitates obtained from Mac-1-HEK293 cells, similar quantities of the Mac-1-CD47 complex can be immunoprecipitated from suspended and adherent cells (Fig. 2, D and E). Since Mac-1-HEK293 cells were used in subsequent structure-function analyses, we characterized these cells in greater detail. In particular, considering CD47 was shown to interact with β_1_ integrins (18,19), and several β_1_ integrins, including α_5_β_1_, α_4_β_1_, α_3_β_1_, α_2_β_1_, and α_v_β_1_, are expressed on the surface of Mac-1-HEK293 cells (36), we determined whether CD47 remained in complexes with β_1_ integrins after precipitation with anti-α_M_. We found that the anti-CD47 mAb could pull down almost all CD47 in a complex with integrin(s) during the initial round of immunoprecipitation (Fig. 2F; denoted 1 IP). Only a negligible amount of CD47 was present in the immunoprecipitates after the second round of precipitation of the supernatant with ani-α_M_ mAb, even though a substantial amount of Mac-1 remained in the lysate (Fig. 2F; denoted 2 IP). Analyses of the immune complexes obtained from the supernatant after the second round of precipitation using anti-β_1_ mAb showed only traces of CD47, suggesting that on the surface of Mac-1-HEK293 cells, the majority of CD47 was in complex with Mac-1 (Fig. 2F; denoted 3 IP).

**Figure 2.**
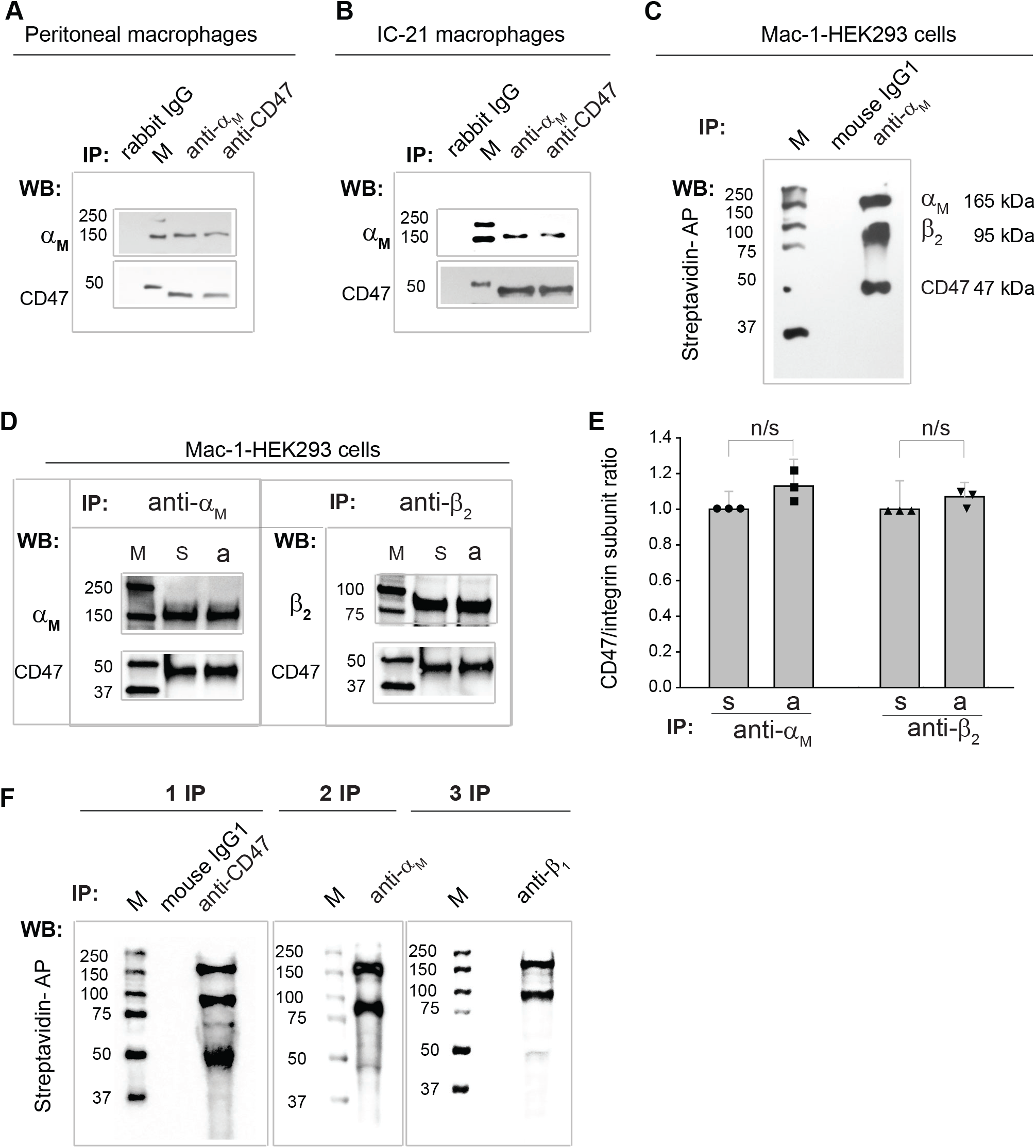
Association of CD47 with Mac-1 on the surface of various Mac-1-expressing cells probed by immunoprecipitation analyses. **(A)** Purified inflammatory peritoneal mouse macrophages were immunoprecipitated with anti-α_M_ rabbit polyclonal antibody or rabbit polyclonal anti-CD47 antibody, and blots were analyzed with rabbit polyclonal antibody against the α_M_ or CD47. Blots of the total cell lysates of WT and CD47-deficient macrophages using rabbit polyclonal antibody against the α_M_ or CD47are shown in the left panel. M, molecular weight markers. The molecular weight of the α_M_ (165 kDa) and β_2_ (95 kDa) integrin subunits and CD47 (47 kDa) are indicated on the right of the panel. **(B)** Murine IC-21 macrophages were immunoprecipitated with anti-α_M_ rabbit polyclonal antibody or rabbit polyclonal anti-CD47 antibody, and blots were analyzed with rabbit polyclonal antibody against the α_M_ or CD47. **(C)** Biotinylated Mac-1-HEK293 cells were lysed and immunoprecipitated with mAb 44a against the α_M_ subunit or isotype control IgG1. Blots were disclosed with streptavidin-alkaline phosphatase (AP). **(D)** Suspended (denoted “s”) or adherent (denoted “a”) Mac-1-HEK293 cells were lysed and immunoprecipitated with anti-α_M_ mAb 44a or anti-β_2_ mAb IB4. Blots were analyzed with anti-α_M_, anti-β_2_, and anti-CD47 antibodies. **(E)** The ratios of CD47 to the α_M_ and β_2_ integrin subunits in the immunoprecipitates from suspended and adherent cells were determined from the densitometry analyses of blots. The ratio of CD47 to each integrin subunit in suspended cells was taken as 1.0. **(F)** Lysates of biotinylated Mac-1-HEK293 cells were immunoprecipitated with anti-CD47 mAb B6H12; then immunoprecipitates were subjected to Western blotting probed with streptavidin-AP (*left panel;* 1 IP). After the first round of immunoprecipitation, the supernatant was immunoprecipitated with anti-α_M_ mAb 44a (*middle panel;* 2 IP). The third round of immunoprecipitation (3 IP) was performed using anti-β_1_ mAb (*right panel*).

### The α_M_ and β_2_ integrin subunits are involved in the interaction with CD47

To determine if the α_M_ or β_2_ subunits of Mac-1 could be implicated in CD47 binding, we generated HEK293 cells expressing individual α_M_ and β_2_ subunits (Fig. 3A) and analyzed them by immunoprecipitation. Also, cells expressing the form of Mac-1 in which the α_M_ I-domain of the α_M_-subunit was deleted were examined. Lysates of α_M_-HEK293 and β_2_-HEK293 cells were incubated with anti-α_M_ mAb 44a or anti-β_2_ mAb IB4 against the α_M_ or β_2_ integrin subunits, respectively, and the immune complexes were analyzed by Western blotting using anti-CD47 mAb (Fig. 3B). The mAbs were able to pull down CD47 from both types of cells suggesting that CD47 can form complexes with both integrin subunits. However, the amount of CD47 in complexes with each integrin subunit was different. As determined by densitometry, while the ratio of CD47 to the α_M_ subunit obtained from α_M_-HEK293 cells was ~2-fold lower than that in immunoprecipitates recovered from cells expressing the whole integrin, the ratio of CD47 to the β_2_ integrin subunit in the immunoprecipitates obtained from β_2_-HEK293 cells was ~1.6-fold greater, suggesting the difference in the strength of complexes (Fig. 3C). Deletion of the α_M_I domain in the I-less integrin did not affect the complex between Mac-1 and CD47, suggesting that this domain is not involved in the CD47 binding (Fig. 3, B and C). Interestingly, the I-less integrin, like the free β2 subunit, bound ~1.5 more CD47 than the whole integrin, which could be tentatively explained by the conformational change in the integrin caused by the deletion of the α_M_I-domain.

**Figure 3.**
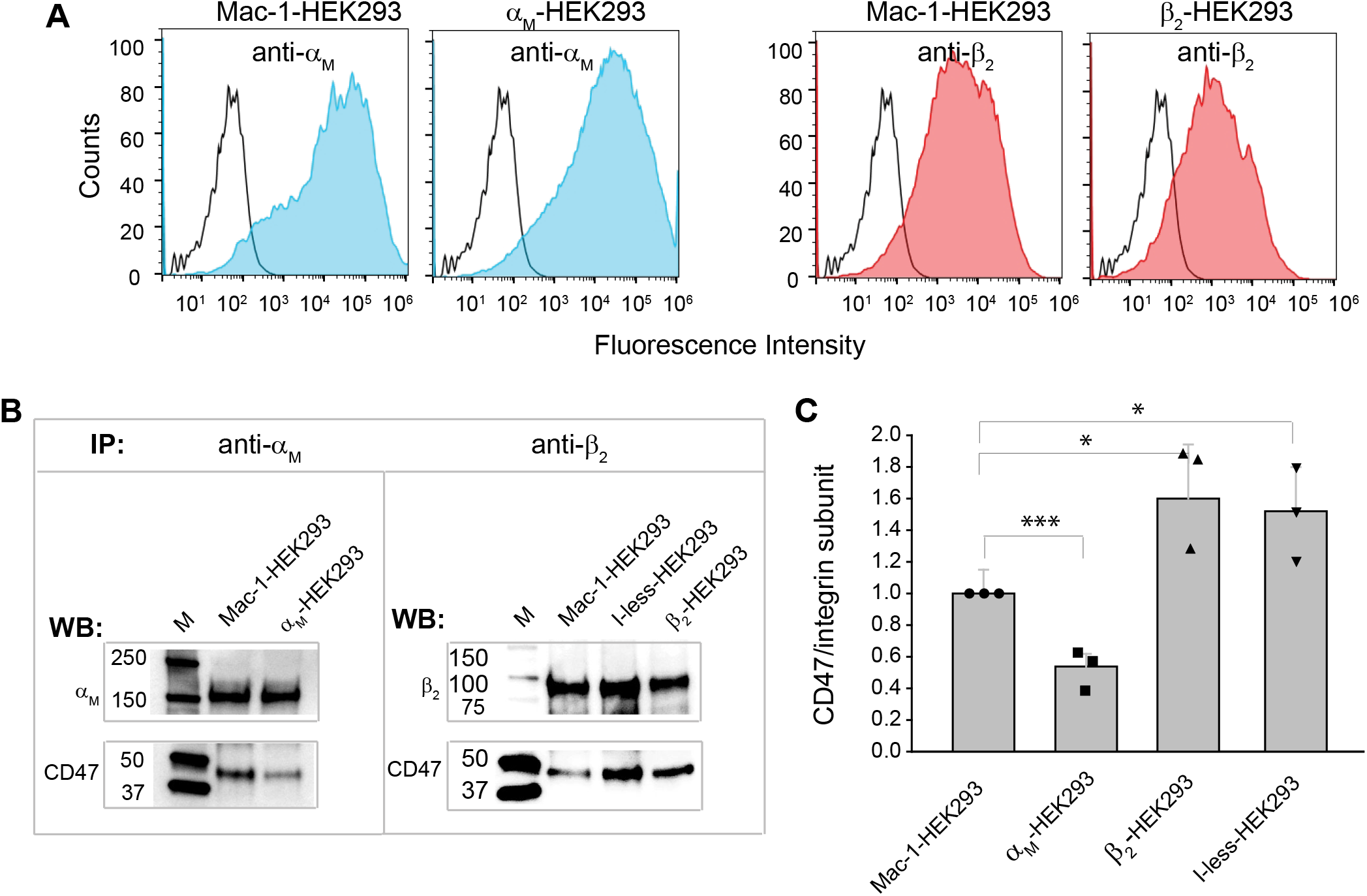
CD47 interacts with both α_M_ and β_2_ subunits of integrin Mac-1. **(A)** Mac-1-HEK293 cells and HEK293 cells expressing individual α_M_ and β_2_ subunits were analyzed by flow cytometry analysis using anti-α_M_ mAb 44a or anti-β_2_ mAb IB4 followed by Alexa Fluor 488-conjugated secondary antibody. Blue histograms indicate cells stained with the primary and secondary antibodies, and open histograms show cells stained with the secondary antibodies (control). **(B)** Lysates of Mac-1-HEK293 cells and cells expressing individual α_M_ and β_2_ integrin subunits were immunoprecipitated with mAb 44a and mAb IB4 against α_M_ and β_2_ integrin subunits, respectively, and immunoprecipitates were analyzed by Western blotting using anti-CD47 mAb. HEK293 cells expressing the I-less form of the α_M_ subunit of Mac-1 were immunoprecipitated with mAb IB4. **(C)** The ratios of CD47 to integrin subunits were determined from densitometry analyses. The ratio of CD47 to either α_M_ or β_2_ subunit in the immune complexes from Mac-1-HEK293 cells was taken as 1.0.

### Activation of Mac-1 increases the amount of CD47 recovered in complexes with Mac-1

The increased reactivity of CD47 with the free β_2_ integrin subunit, which is known to exist in the extended conformation when expressed alone (37), suggested that the binding site for CD47 in the whole receptor may be partially shielded due to the bent conformation of integrin and becomes exposed in the extended integrin. To investigate this possibility, we determined the amount of CD47 in complexes with integrin immunoprecipitated from Mac-1-HEK293 cells treated with Mn^2+^, PMA, and mAb MEM48. It is known that the activation of β_2_ integrins with these reagents converts integrins from the bent to the extended conformation that can be detected by mAb KIM127. The mAb KIM127 recognizes Gly504, Leu506, and Tyr508, the residues at the end of EGF-2 (37) shielded in the interface with the PSI domain in the bent integrin conformation and became exposed in the extended conformation (38,39). Initial FACS analyses showed that 12 ± 2% of Mac-1 on the surface of nonactivated Mac-1-HEK293 cells bound KIM127, indicating that a portion of molecules was already in the extended conformation (Fig. 4A). The treatment of cells with 100 nM PMA and 1 mM Mn^2+^ resulted in a ~1.8 and ~2.6-fold increase in the number of cells presenting Mac-1 in the extended conformation (Fig. 4A). In addition, activation of cells with mAb MEM48 followed by Alexa Fluor 488-labeled KIM127 showed a ~2-fold increase of cells presenting extended receptors (Fig. 4A, *bottom panel).* In agreement with the transition of Mac-1 into an extended conformation, adhesion of Mn^2+^-activated cells to fibrinogen was increased (Fig. S2). Mac-1-CD47 complexes immunoprecipitated from cells activated with PMA and Mn^2+^ contained more CD47 than nonactivated cells (Fig. 4, B and C). The activation of cells with MEM48 resulted in an even higher amount of CD47 complexed with Mac-1 (Fig. 4, B and C).

**Figure 4.**
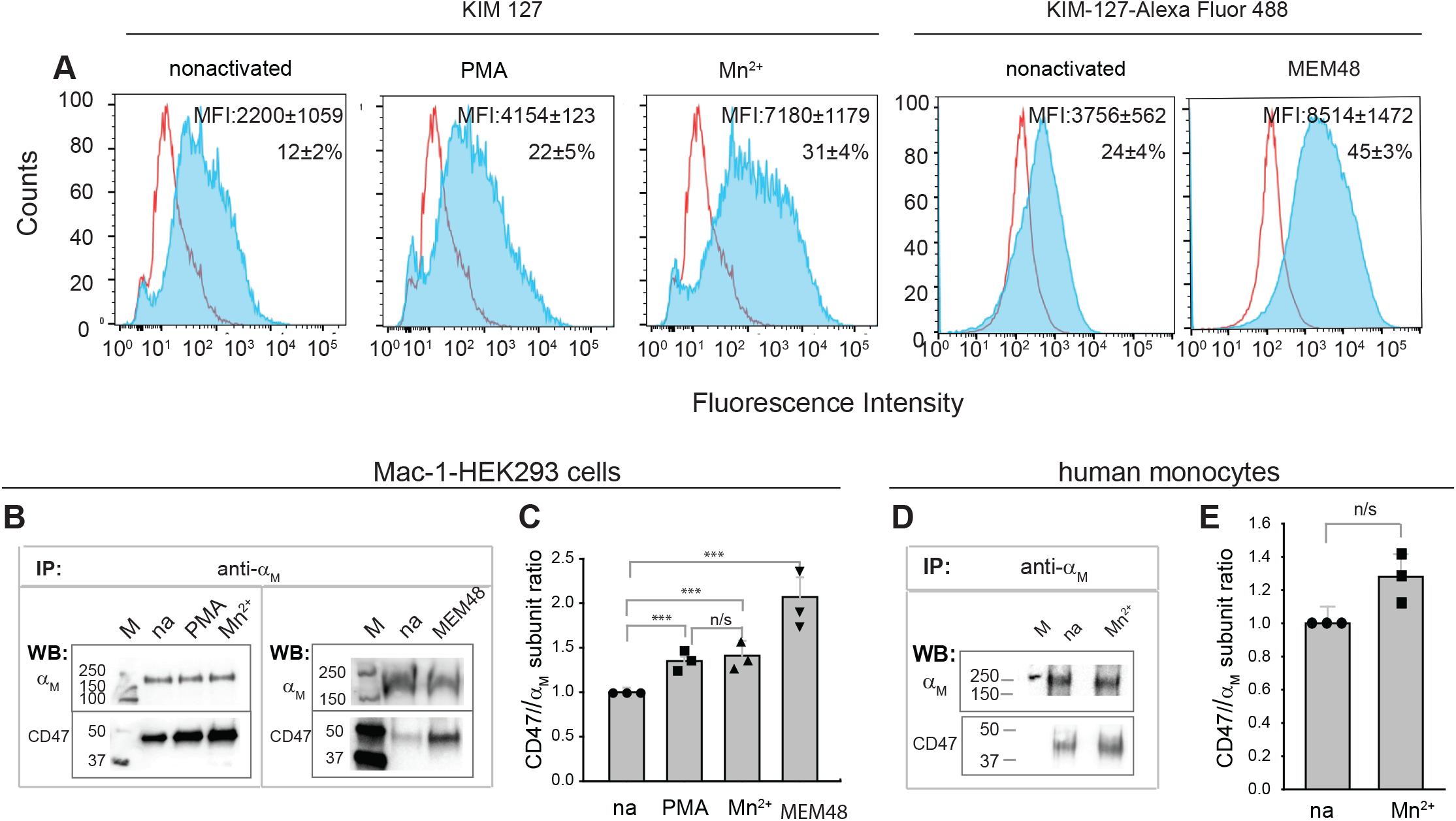
Activation of Mac-1-HEK293 cells results in increased CD47 recovered in complexes with Mac-1. **(A)** The Mn^2+^- and PMA-induced conversion of Mac-1 into the extended conformation was detected by FACS analysis using the reporter mAb KIM127 (*upper panel*). Blue histograms indicate cells incubated with KIM127 and Alexa Fluor 488-conjugated secondary antibody antibodies, and open histograms show cells incubated with the secondary antibodies (control). To determine the epitope expression for mAb KIM127 after activating mAb MEM48, mAb KIM127 was conjugated to Alexa Fluor 488 (*bottom panel*). Blue histograms are cells stained with Alexa Fluor 488-conjugated KIM127, and open histograms show unstained cells (control). The number of KIM127 positive cells is a percent of cells expressing Mac-1, determined in each experiment with anti-α_M_ mAb 44a and typically constituted 90 ±3% of a total cell population. Histograms are representative of three separate experiments. Data are means ± SE. **(B)** Lysates of Mac-1-HEK293 cells activated with 100 nM PMA, 1 mM Mn^2+^ and MEM48 (5 μg/ml) were immunoprecipitated with anti-α_M_ mAb 44a, and the precipitates were analyzed by Western blotting using antibodies against α_M_ and CD47. **(C)** The ratio of CD47 to the α_M_ subunit was determined from densitometry analyses. The ratio of CD47 to α_M_ in the immune complex obtained from nonactivated Mac-1-HEK293 cells was assigned 1.0. **(D)** Lysates of nonactivated and Mn^2+^-stimulated human monocytes were immunoprecipitated with mAb 44a and analyzed by Western blotting with antibodies against α_M_ and CD47. **(E)** The ratio of CD47 to α_M_ in the immune complex obtained from nonactivated cells was assigned 1.0. Data shown are means ± SE from three individual experiments.

The presence of a portion of Mac-1 molecules in the extended state on the surface of nonactivated Mac-1-HEK293 cells raised the question of whether coprecipitation of CD47 with Mac-1 was a property of the extended integrins and that bent integrins were not able to bind CD47. Similarly, immunoprecipitation of Mac-1-CD47 complex from cultured IC-21 murine macrophages and macrophages isolated from the inflamed peritoneum might have been due to the activated state of these cells. We performed immunoprecipitation analysis using freshly isolated human monocytes to examine whether CD47 does not bind the nonactivated integrin on resting cells. Fig. 4, D and E show that mAb 44a precipitated Mac-1 in complex with CD47, and activation of monocytes with Mn^2+^ increased the amount of CD47, suggesting that in resting monocytes, CD47 is prebound to Mac-1 and cellular activation, through the extension of the integrin, strengthens the complex.

To examine further whether CD47 regulates the integrin conformation, we disrupted CD47 gene expression in Mac-1-HEK293 cells using the CRISPR/Cas9 technology. Human cells were used for these experiments because conformation-sensitive antibodies against mouse β_2_ integrins are unavailable. FACS analyses of selected cells lacking CD47 (Fig. 5A) showed unaltered Mac-1 expression (Fig. 5B). Consistent with adhesion studies of CD47-deficient mouse macrophages, adhesion of CD47-deficient Mac-1-HEK293 cells was significantly reduced (Fig. S3). Furthermore, considerably fewer integrins on the surface of CD47-deficient cells treated with Mn^2+^ expressed the KIM127 epitope, suggesting the failure of Mac-1 to convert into the extended conformation (Fig. 5, C and D). The results indicate the Mac-1-CD47 interaction is strengthened after converting integrin from the bent to the extended conformation and further suggest that CD47 is required to stabilize the extended integrin conformation.

**Figure 5.**
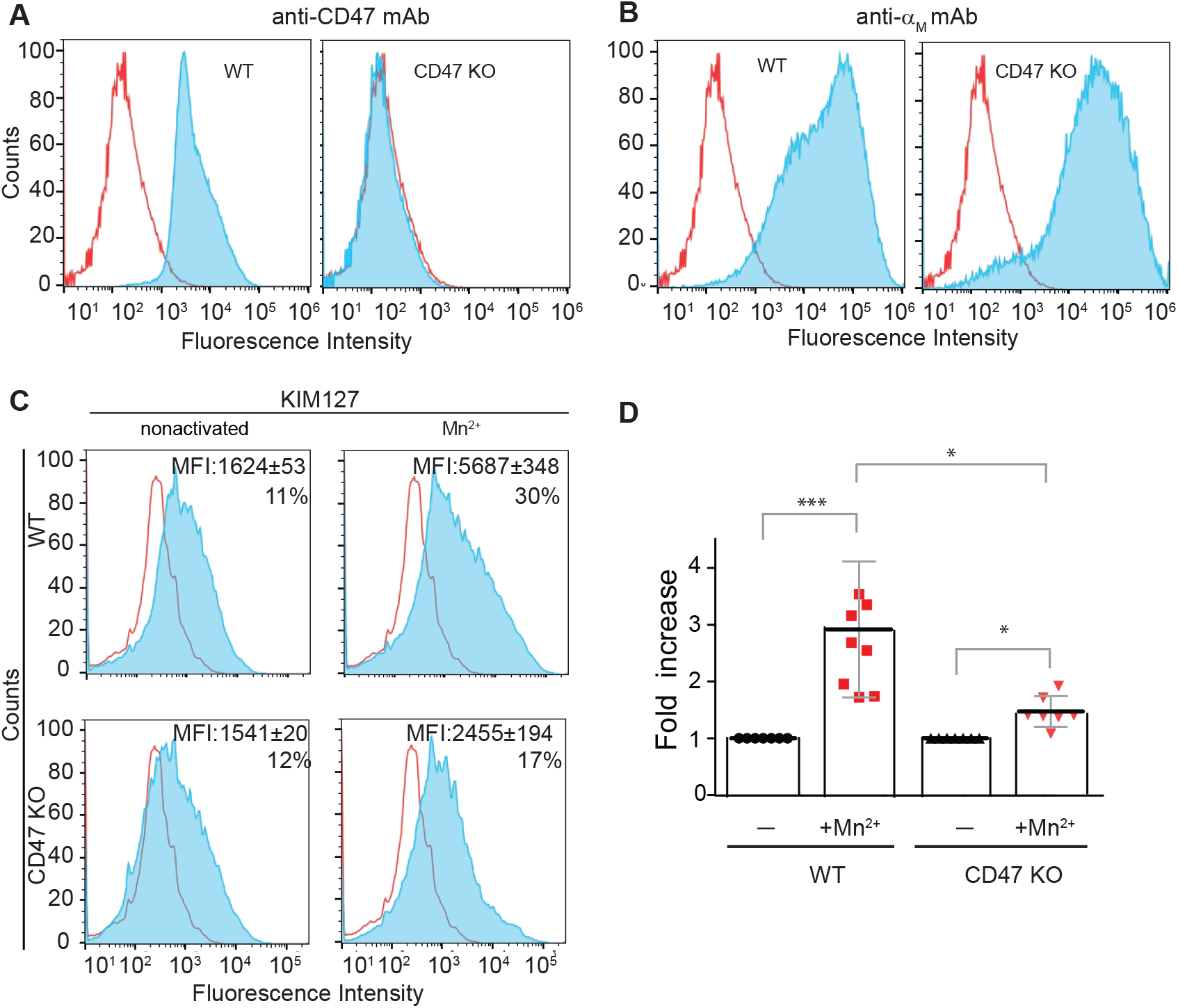
CD47 deficiency in Mac-1-HEK293 cells results in reduced expression of the KIM127 epitope in response to Mn^2+^. Expression of CD47 in WT Mac-1-HEK293 cells **(A)** and Mac-1-HEK293 cells in which CD47 gene expression was disrupted using the CRISPR/Cas9 technology **(B)** was analyzed by FACS with anti-CD47 mAb B6H12 and anti-α_M_ mAb 44a. Blue histograms indicate cells stained with the primary and secondary antibodies, and open histograms show cells stained with the secondary antibodies (control). **(C)** WT and CD47-deficient Mac-1-HEK293 cells were treated with 1 mM Mn^2+^ and the conversion of Mac-1 into the extended conformation was detected by FACS using mAb KIM127 (blue histograms). Representative histograms of three separate experiments are shown. The numbers indicate the percentage of positive cells. **(D)** The binding of mAb KIM127 after treating WT and CD47-deficient Mac-1-HEK293 cells with Mn^2+^ is shown as a fold increase in the percent of positive cells compared to untreated cells.

### The IgV domain of CD47 binds to Mac-1

To localize the binding site for Mac-1 in CD47, we tested the idea that the single extracellular IgV domain of CD47 is involved in the interaction with Mac-1. The recombinant IgV domain was expressed in *E.coli* as a fusion protein with GST (Fig. 6A), and its ability to compete with the CD47-Mac-1 complex formation was examined. As shown in Fig. 6B and 6C, incubation of Mac-1-HEK293 cells with increasing concentrations of IgV-GST resulted in the dose-dependent decrease of CD47 that can be recovered in immune complexes, suggesting that recombinant protein can displace the IgV domain of CD47 in complex with Mac-1. Purified control GST was not effective (Fig. 6B, C). Recombinant GST-IgV was not detected in complex with Mac-1, but this may result from a weaker affinity of isolated IgV compared to the IgV domain in the context of the whole molecule.

**Figure 6.**
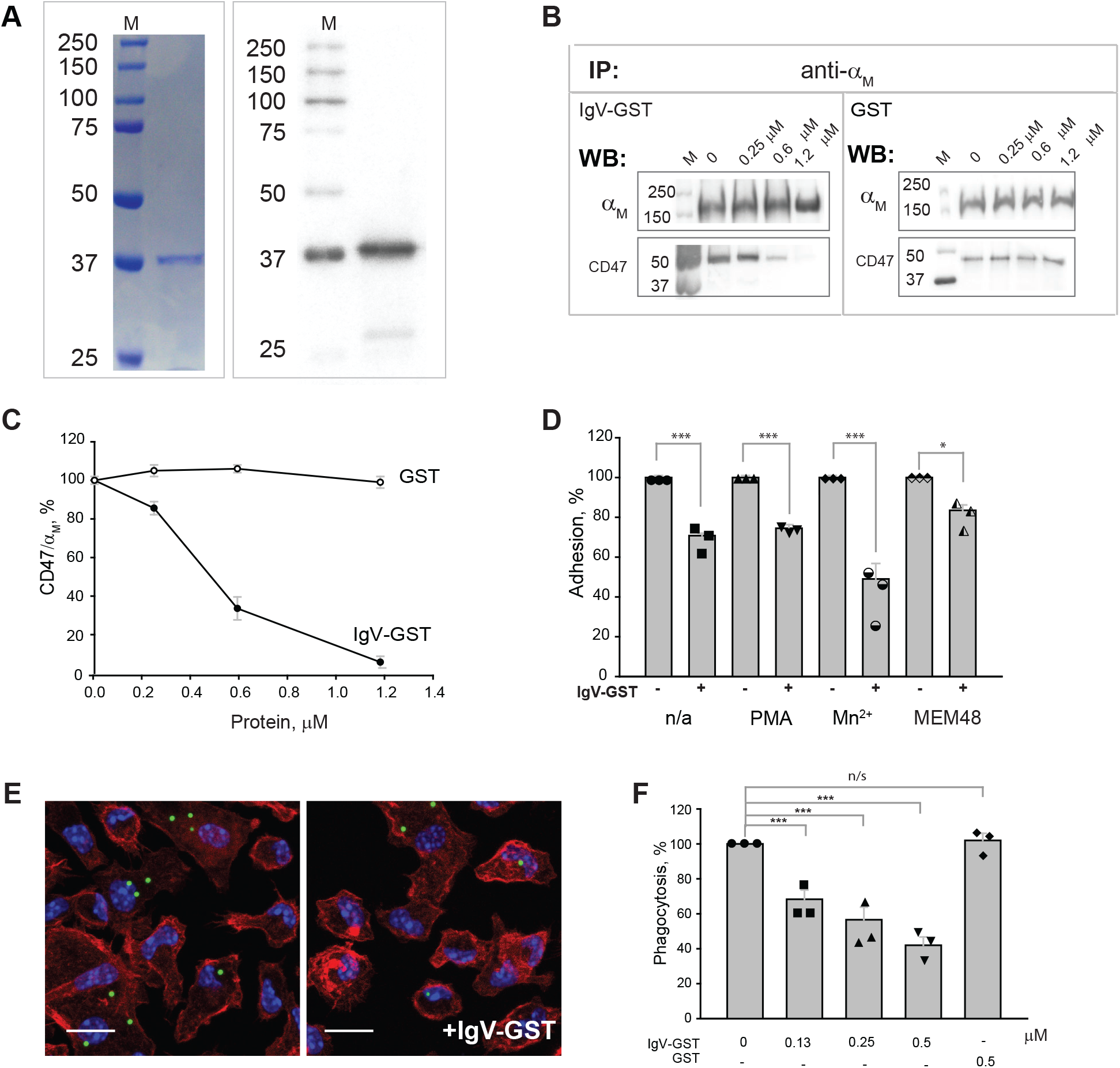
The IgV domain of CD47 binds Mac-1. **(A)** Recombinant IgV domain of CD47 fused with GST was purified from soluble fractions of *E.coli* lysates by affinity chromatography using glutathione agarose and characterized by SDS PAGE *(left panel)* and Western blotting *(right panel).* (B) Mac-1-HEK293 cells were incubated with different concentrations of IgV-GST *(right panel)* or GST *(right panel).* CD47 in complex with Mac-1 was determined after immunoprecipitation of lysates using mAb 44a, followed by Western blotting with anti-CD47 antibody. **(C)** Effect of IgV-GST and GST on complex formation between CD47 and Mac-1 determined from immunoprecipitation analyses shown in (B). The CD47/α_M_ ratio in the absence of IgV-CD47 was assigned a value of 100%. **(D)** Effect of IgV-GST on adhesion of nonactivated and activated Mac-1-HEK293 cells to fibrinogen. Calcein-labeled nonactivated and PMA, Mn^2+^, or mAb MEM48-stimulated cells were preincubated with 1.2 μM IgV-GST for 15 minutes at 22° C. Adhesion in the absence of IgV-GST was assigned a value of 100%. Data shown are mean ± SE from 3 separate experiments. *p≤ 0.1; ***p≤ 0.001 compared with control adhesion in the absence of IgV-GST. **(E)** The effect of IgV-GST on phagocytosis of LL-37-coated latex beads by peritoneal macrophages isolated from WT mice. Adherent macrophages were incubated with beads in the absence or presence of different concentrations of IgV-GST for 30 min at 37 °C, and non-phagocytosed beads were removed by washing. **(F)** Phagocytosis was quantified as a percentage of uptake of LL-37-coated beads in the absence of IgV-GST. Phagocytosis was determined from ten fields of fluorescent images. Data shown are means ± SE from three separate experiments.

To further demonstrate the importance of the IgV domain of CD47 in binding to Mac-1, we examined the effect of recombinant IgV on selected Mac-1-mediated functional responses. The preincubation of Mac-1-HEK293 cells with 1.2 μM IgV-GST resulted in a partial but significant decrease in adhesion of both nonactivated and activated cells to fibrinogen (Fig. 6D). The recombinant IgV more effectively inhibited adhesion of cells activated with Mn^2+^ (51 ± 7%) than cells activated with PMA and mAb MEM48 (25 ± 0.9% and 18± 3%, respectively). As a control, GST was inactive. These results demonstrate that recombinant IgV can compete with IgV in membrane-bound CD47. However, since isolated IgV reduces cell adhesion only partially, the transmembrane or cytoplasmic portions of CD47 may be required for its function. To examine the effect of recombinant IgV on opsonin-mediated phagocytosis, adherent mouse macrophages were incubated with IgV-GST for 30 min, and then LL-37-coated beads were added. Recombinant IgV inhibited phagocytosis of opsonized beads in a dose-dependent manner with the IC_50_ value of 0.36 ± 0.04 μM (Fig. 6, E and F). Incubation of cells with GST did not inhibit phagocytosis. These observations indicate that the Mac-1 binding site likely resides within the IgV domain of CD47.

### The membrane-proximal β_2_ EGF-3-4 and α_M_ calf-1-2 domains of Mac-1 interact with the IgV domain of CD47

The dimensions of the IgV domain of CD47 plus a short linker connecting its C-terminal end and the first transmembrane segment suggest that IgV may protrude from the plasma membrane at a distance of ~5 nm (40). In addition, a long-range disulfide bond between Cys15 in the IgV domain and Cys245 in the extracellular loop between the fourth and fifth transmembrane helices (41) keeps IgV close to the plasma membrane (40). Such positioning may juxtapose the IgV domain of CD47 with the membrane-proximal region of Mac-1 encompassing the β tail (βT), EGF-4, and the C-terminal part of EGF-3 domains of the β_2_ subunit, whose lengths are 2.5, 2.0 and 2.1 nm, respectively (Fig. 7A). In the bent conformation, EGF-3 and EGF-4 form the tight interface and are partially shielded by the α_M_ β-propeller and β_2_ hybrid domains, which may explain the greater amount of CD47 in complex with Mac-1 after the conversion of integrin into the extended conformations caused by activating stimuli. Furthermore, since CD47 was immunoprecipitated with the α_M_ subunit from α_M_-HEK293 cells, the IgV domain can also potentially interact with the membrane-proximal calf-2 and the C-terminal portion of calf-1, whose lengths are ~4 nm each. Based on these considerations, we prepared recombinant fragments derived from the part of the β_2_ subunit spanning EGF-3 through βT (Fig. 7, B and C) and the calf-1 and calf-2 domains of the α_M_ subunit and examined their interaction with CD47-IgV. The proteins were prepared with a 6xHis tag, and their binding to immobilized CD47-IgV freed from its fusion part was analyzed using a solid-phase binding assay. Figure 7D shows that fragments containing EGF-3 and EGF-4, including EGF-3-4-βT and EGF-4-βT, as well as isolated EGF-3 and EGF-4 bound CD47-IgV in a dose-dependent manner. The binding of the βT fragment was negligible. To substantiate the involvement of the β_2_ subunit-derived fragments in CD47 binding, we examined the ability of these proteins to inhibit Mac-1-CD47 complex formation on the surface of Mac-1-HEK293 cells. As shown in Fig. 7E and 7F, EGF-3-4-βT inhibited complex formation between Mac-1 and CD47 in a dose-dependent manner. The βT fragment was not active. The interaction of recombinant calf-1 and calf-2 fragments with CD47-IgV was also detected (Fig. 7G). However, when tested at the same concentration (5 μM), calf-1 and calf-2 were less active than the EGF-3-4-βT fragment. The binding of the control recombinant IgV domain derived from mouse SIRPα was insignificant (Fig. 7G).

**Figure 7.**
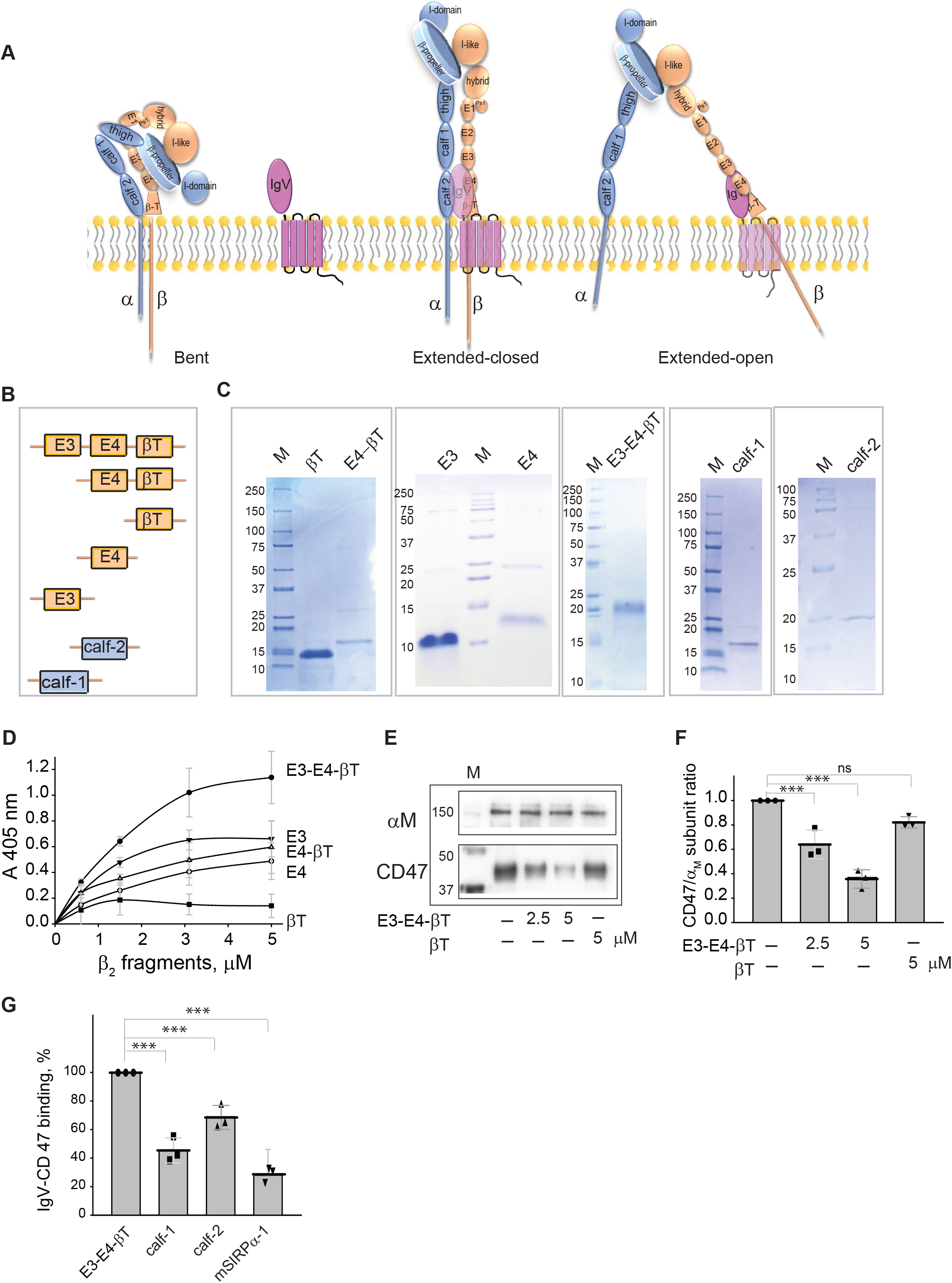
Localization of the binding site for CD47-IgV in Mac-1. **(A)** Schematic representation of the integrin Mac-1 in the bent, extended-closed, and extended-open conformations. CD47 is prebound to the nonactivated bent integrin (not shown). CD47 is arbitrarily shown as bound to both integrin subunits in the extended-closed conformation and the β_2_ subunit in the extended-open conformation. **(B)** Schematic of recombinant proteins used in solid-phase binding assay and immunoprecipitation experiments. E3, E4, and βT denote EGF-3, EGF-4, and β-tail fragments derived from the β_2_ subunit, and calf-1 and calf-2 are derived from the α_M_ subunit. **(C)** His-tagged recombinant proteins were purified from soluble fractions of *E.coli* lysates by affinity chromatography using Ni-NTA-agarose column and characterized by SDS-PAGE. M, markers. **(D)** Binding of recombinant fragments to CD47-IgV. The wells of microtiter plates were coated with free CD47-IgV (2 μg/ml), and different concentrations of recombinant fragments were added to the wells for 3 h at 37 °C. Bound proteins were detected using an anti-His mAb followed by an AP-conjugated secondary antibody. Data are the mean ± SE from 3 separate experiments. **(E)** Mac-1-HEK293 cells were incubated without or with different concentrations of E3-E4-βT (2.5 and 5 μM) or βT (5 μM) and the CD47-Mac-1 complex formation was determined after immunoprecipitation of lysates using mAb 44a followed by Western blotting. **(F)** CD47 in complex with Mac-1 in immunoprecipitates was determined from densitometry analyses. The ratio of CD47 to the α_M_ subunit in immune complexes in the absence of inhibitors was taken as 1.0. **(G)** The binding of calf-1 and calf-2 fragments to immobilized IgV-CD47 was compared with E3-E4-βT. Recombinant proteins were tested at equimolar concentrations (5 μM). The recombinant IgV domain of mouse SIRPAα (mSIRPα-1) was used as a negative control. The binding of E3-E4-βT to IgV-CD47 was taken as 1.0. Data shown are mean ± SE from 3 separate experiments.

## DISCUSSION

CD47 plays an essential role in regulating functions of integrins expressed on various cells, including leukocytes, where it was found in complexes with α_v_β_3_ (VLA-3) and α_L_β_2_ (LFA-1) (5–7,20). The present study demonstrates that CD47 forms a complex with and regulates adhesive functions of the major myeloid cell-specific integrin Mac-1 and clarifies the basis for its modulating effect. In particular, the lack of CD47 in mouse macrophages strongly impairs adhesion, spreading, migration, phagocytosis, and macrophage fusion. Based on co-immunoprecipitation analyses, previous studies suggested a physical link between CD47 and β_1_ and β_3_ integrins (5–7,15,17,18), (19). However, only a single study detected the direct interaction of CD47 with integrin α_IIIb_β_3_, albeit in the presence of the thrombospondin-binding peptide 41NK (16). Localizing the regions in CD47 and Mac-1 involved in their association, our study provides the first evidence for the direct interaction between these molecules. We show the interaction between the IgV domain of CD47 and the membrane-proximal EGF-3 and EGF-4 domains of the β_2_ integrin subunit and calf-1 and calf-2 of the α_M_ subunit is responsible for the CD47-Mac-1 complex formation.

Our data demonstrate that CD47 is prebound to Mac-1, as evidenced by coprecipitation of CD47 with Mac-1 by anti-Mac-1 and anti-CD47 antibodies from resting monocytes and nonactivated Mac-1-expressing HEK293 cells. It is well established that on the surface of resting cells, integrins are mostly maintained in a default inactive state exhibiting a bent conformation (38,39,42,43). In this conformation, the integrin headpiece formed by the head and upper leg domains of the α and β subunits closely contacts the lower legs (Fig. 7A). The bends occur at the knees between the thigh and calf-1 domains in the α subunit and EGF-1 and EGF-2 in the β subunit. The bent conformation is also stabilized by the association between the cytoplasmic parts of the integrin α and β subunits (37). The release of the interface between the headpiece and lower legs at the α- and β-knees converts the integrin into an extended conformation (Fig. 7B) (38,39). This conformation with low ligand affinity has been termed “extended-closed” (26). In addition to the extension at the α and β knees, the headpiece opening resulting from the conformational changes in the βI-like domain in the β subunit and the adjacent hybrid domain forces the latter to swing away, leading to leg separation (Fig. 7C) (26). This integrin conformation is known as “extended-open.” Structural studies on several integrins have demonstrated a direct relationship between the extended-open conformation and high affinity for ligands (rev. in (44)). Previous studies showed that extracellular activation of integrins by Mn^2+^, as well as inside-out activation of integrins stimulated by protein kinase C, induce an extended integrin conformation reported by the conformation-sensitive mAb KIM127 (45). Our findings that the amount of CD47 in complexes with Mac-1 was increased after integrin activation by Mn^2+^ and PMA suggest that the CD47 binding site in Mac-1 is partially shielded in the bent conformation and becomes available in the extended conformations. This proposal also agrees with the higher amount of CD47 in the complex with the free β_2_ subunit (Fig. 3), which is known to exhibit extended conformation when expressed alone (37). We observed an even greater effect of activating mAb MEM48, the epitope for which was mapped to the interface between the EGF-3 and EGF-4 (37). However, the reason for this is unclear.

We showed that the binding site for Mac-1 in CD47 resides in the IgV domain of CD47, and the complementary binding sites in Mac-1 are present in the EGF-3-4 and calf-1-2 domains of the β_2_ and α_M_ subunits, respectively. The location of these sites in the membrane-proximal parts of integrin legs is consistent with close adjacency of CD47-IgV to the plasma membrane in the context of the whole CD47 (40). In the three-dimensional structure of αxβ_2_ (CD11c/CD18), a sister integrin highly homologous to Mac-1, some portions of EGF-3-4 and calf-1-2 are exposed in the bent integrin on resting cells and may be available for CD47-IgV binding (39). However, significant portions of both regions are shielded in the bent conformation by the α subunit β-propeller and β_2_ hybrid domain and by close apposition between EGF-3-4-βT and calf-1-2. The elongation of integrin in the extended-close conformation may expose some amino acid sequences and make them available for CD47-IgV interaction. Furthermore, the separation of legs in the extended-open conformation would disrupt the interface between the inner faces of EGF-3-4 and calf-1-2 and unmask additional sites for the interaction with CD47-IgV. The increase in CD47 in complex with Mac-1 after activation with PMA and Mn^2+^ is consistent with such an explanation; however, further studies of CD47-Mac-1 binding sequences are required to substantiate the proposed model.

What is the possible role of CD47 in the complex with Mac-1? The three-dimensional structure of the IgV domain of CD47 shows that its dimensions of ~5 nm x 3.2 nm (40) could theoretically permit the interaction with both EGF-3-4 and calf-1-2 that are adjacent in the bent and extended-closed conformations. However, the separation of legs in the extended-open conformation at a distance of ~12 nm (44) should preclude the simultaneous engagement of two integrin legs by a single IgV domain. A significantly higher amount of CD47 recovered from cells expressing a free β_2_ subunit than cells expressing the α_M_ subunit or the whole receptor suggests that the β_2_ subunit has a higher affinity for CD47. Based on this finding, it seems plausible to assume that the EGF-3-4 may associate with CD47-IgV after the leg separation. However, the possibility that the α and β subunits interact with the IgV domains on different CD47 molecules cannot be ruled out. The EM studies of integrins α_x_β_2_, α_L_β_2_, α_v_β_3_, and α_5_β_1_ demonstrated significant variability in the lower β leg orientation below the PSI-EGF1 domains (26,38,44). Furthermore, the ten different β_2_ leg structures of α_x_β_2_ were observed in three crystal lattices (39). These findings have led to a proposal that the β legs are highly flexible (26). The IgV domain of CD47, through its disulfide bond to the extracellular loop connecting the fourth and fifth transmembrane segments (40), seems to be constrained in the defined orientation in the membrane-bound CD47. Therefore, it is tempting to hypothesize that the IgV may retard the lateral motion of the β_2_ leg in the plasma membrane plane and stabilize the extended-open conformation of Mac-1 and other β_2_ integrins required for the high-affinity ligand binding. Such stabilization may also be necessary for coupling the β_2_ leg with the intracellular signaling machinery, as illustrated by the markedly weak spreading of CD47-deficient macrophages (Fig. 1, C and E).

We observed a striking defect in the epitope exposure for the activation reporter mAb KIM127 in CD47-deficient Mac-1-HEK293 cells after their treatment with Mn^2+^. These data are in line with previous studies by Azcutia et al. (20). These investigators showed reduced binding of mAbs KIM127 and 24 to α_L_β_2_ after activation of CD47-deficient T cells with Mn^2+^ and proposed that CD47, through its association with integrin, directly or indirectly facilitates transition into or stabilizes the activated form of α_L_β_2_. A defect in the binding of the anti-β_1_ activation reporter mAb HUTS21 in CD47-deficient T cells has also been detected (20). The reduced exposure of epitopes for the activation reporter mAbs in CD47-deficient cells implies that either a portion of integrins on the surface of activated cells remained in the bent conformation or integrins adopted a state intermediate between the bent and extended conformations. Yet another possibility is that some integrins transitioned from the extended conformation back into the bent conformation. In this regard, Tang et al. reported the reversion of Mg/EGTA-activated α_L_β_2_ that was reactive with KIM127 to its resting nonreactive bent conformation after depletion of activating cation by EDTA (46). These different possibilities are theoretically not mutually exclusive and align with the idea that CD47 exerts the stabilizing effect on the extended conformation of integrin.

Based on the negative stain EM studies with various activating and blocking mAbs, it has been proposed that the β_1_, β_2_, and β_3_ subfamilies of integrins have similar overall global conformational states, i.e., bent-closed, extended-closed, and extended-open (26,47). Since the single CD47 molecule regulates functions of all three integrin subfamilies expressed on various cells, the common mechanism involving the interaction of CD47-IgV with the EGF-3-4 and calf-1-2 may underlie the CD47 function. Although the EGF-3-4 domains of β_1_, β_2_, and β_3_ integrin subunits exhibit moderate homology (except for conservative cysteines), all three share a high degree of homology in the segment corresponding to ^568^CSGRGRC^574^ in EGF-4 of the β_2_ subunit and some amino acid residues. The contribution of EGF-3-4 and calf-1-2 domains of the β_1_ and β_3_ subunits to the interaction with CD47-IgV remains to be determined.

In addition to regulating the integrin functions, CD47 serves as a counter-receptor for signal-regulatory protein-alpha (SIRPα), which is abundantly expressed in macrophages (48,49). In a number of homeostatic and inflammatory processes, the CD47-SIRPα interaction sends the inhibitory “do not eat me” signal in macrophages, preventing the destruction of normal host cells (50–53). Based on this phenomenon, CD47 has been implicated as a “marker of self” (2). Since many hematologic and other malignancies demonstrate elevated levels of CD47 (54–56), it has been proposed that CD47 may allow cancer cells to evade phagocytosis-mediated elimination (54). Accordingly, targeting CD47 may be a unique mechanism of action with broad applicability in anti-tumor therapy aimed to augment macrophage-mediated destruction of cancer cells, and consequently, several antibodies that target CD47 have been generated (57,58)Zhang, 2020 #8548}. It is known that the CD47-SIRPα interaction can regulate phagocytosis mediated by two main phagocytic systems on macrophages, Fcγ and complement receptors (59). One of the complement receptors is the integrin Mac-1, also known as the complement receptor 3 (CR3)(60). Our data show that CD47 is essential for the phagocytic function of Mac-1 on macrophages. Therefore, ideally, the antibodies that target CD47 during anti-tumor therapy should inhibit the *trans* CD47-SIRPα interaction but spare the *cis* CD47-Mac-1 complex for its full phagocytic activity. One of the best-characterized anti-CD47 mAbs that showed efficacy in numerous mouse tumor models is mAb B6H12 (61). Although effective in targeting the CD47-SIRPα interaction, this mAb is also known to inhibit many integrin-dependent functions, including phagocytosis (9,11,13,62). The crystal structure of the CD47-B6H12 complex showed that the SIRPα binding site in CD47 overlaps with epitopes recognized by mAb B6H12 (40,63), suggesting that the binding sites in CD47 for integrins and SIRPα may partially or fully overlap. Thus, it will be interesting and important to determine the critical determinants responsible for the interaction of CD47-IgV with the EGF-3-4 and calf-1-2 domains of Mac-1.

In conclusion, our studies provide evidence for the direct interaction between the integrin Mac-1 and CD47 and explain the molecular basis for their association. Significantly, we show that the Mac-1-CD47 interaction is enhanced after cell activation, which may be essential for stabilizing the extended state of integrin required for high-affinity ligand binding involved in many responses of this immunologically important integrin.

## EXPERIMENTAL PROCEDURES

### Reagents

Human fibrinogen and thrombin were obtained from Enzyme Research Laboratories (South Bend, IN). ICAM-1 was obtained from R&D Systems, Inc. (Minneapolis, MN). The cathelicidin peptide LL-37 was synthesized by Peptide 2.0 (Chantilly, VA). The mouse mAb 44a directed against the human α_M_ integrin subunit, the rat mAb M1/70, which recognizes both mouse and human α_M_ integrin subunits, the mouse mAb IB4 directed against the human β_2_ integrin subunit, the mouse mAb KIM127 which recognizes the extended conformation of human β_2_ integrins and the mouse mAb B6H12 directed against human CD47 were purified from conditioned media of hybridoma cells obtained from the American Tissue Culture Collection (Manassas, VA) using protein A agarose. The mouse mAb MEM48 which recognizes the human β_2_ integrin subunit (catalog #CBL158) and the mouse mAb 1965 against the human β_1_ integrin subunit (catalog #MAB1965) were from EMD Millipore (Burlington, MA). The rat PE-conjugated mAb against F4/80 (catalog #12-4801-82) was from Thermo Fisher. The rabbit polyclonal anti-mouse α_M_ antibody (catalog #ab128797) was from Abcam (Cambridge, MA). The mouse anti-human α_M_ mAb (catalog #66519-1-Ig) and the rabbit polyclonal antibody, which recognizes both mouse and human β_1_ integrin subunits (catalog #12594-1-AP), were from Proteintech (Rosemont, IL). The rabbit polyclonal anti-human/mouse CD47 antibody (catalogue #CD47-101AP) was from Fabgenix (Frisco, TX). The rat monoclonal anti-CD47 antibody (catalog #sc-12731) and mouse monoclonal anti-His-tag antibody (catalog #sc-8036) were from Santa Cruz Biotechnology (Santa Cruz, CA). The secondary antibodies, goat anti-mouse IgG (H+L) conjugated to horseradish peroxidase (HRP) (catalog #1706516) and goat anti-rabbit IgG (H+L)-HRP (catalog #1721019), were obtained from Bio-Rad (Hercules, CA). HRP-conjugated goat anti-rat IgG (catalog #62-9520) and Zysorbin-G (catalog #10-1051-1) were from Invitrogen (Carlsbad, CA). The mouse IgG1 (catalog #401407), an isotype control for mAbs 44a and B6H12, and the rat IgG2b (catalog #400621), an isotype control for M1/70 were obtained from BioLegend (San Diego, CA). Alexa Fluor 568-conjugated phalloidin (catalog #A12380), Alexa Fluor 488-conjugated phalloidin (catalog #A12379), Calcein-AM (catalog #C3100MP), Alexa Fluor 488 Antibody Labeling kit (A20181) and fluorescent latex beads (FluoroSpheres, 1 μm) were from Thermo Fisher. Interleukin-4 (catalog #Z02996) was purchased from GenScript (Piscataway, NJ). Reduced glutathione (catalog #G6529) and the protease inhibitor cocktail (Sigma-Aldrich) were from Sigma-Aldrich (St. Louis, MO). Glutathione-agarose was from ThermoFisher Scientific.

### Mice

Wild-type C57BL/6 and CD47^-/-^ *(B6.129S7-Cd47^tm1Fpl^/J)* mice were obtained from The Jackson Laboratory. All procedures were performed under the animal protocols approved by the Institutional Animal Care and Use Committee of Arizona State University. Animals were maintained under constant temperature (22 °C) and humidity on a 12-h light/dark cycle in the Animal Facility of Arizona State University. Eight-to 12-week-old male and female mice were used in all experiments with age- and sex-matched wild-type (WT) and deficient mice selected for side-by-side comparison. Peritonitis in mice was induced by the intraperitoneal injection of 0.5 mL of a 4% Brewer thioglycollate (TG) solution (Sigma-Aldrich, St. Louis, MO).

### Cells

Human embryonic kidney cells (HEK293) stably expressing integrin Mac-1 (Mac-1-HEK293) were previously described (36,64). HEK293 cells stably expressing the α_M_ subunit or transiently expressing the β_2_ subunit of Mac-1 were generated using a pcDNA3.1 vector containing the full-length cDNAs from α_M_ or β_2_ essentially as described (64). HEK293 cells expressing a heterodimer consisting of the modified α_M_ subunit in which α_M_I-domain was deleted and intact β_2_ subunit (denoted I-less-HEK293 cells) were generated as previously described (65). Mac-1-HEK293 cells in which CD47 gene expression was disrupted were generated using CD47 CRISPR/Cas9 KO Plasmid (h) obtained from Santa Cruz Biotechnology (catalog #sc-400508). According to the manufacturer’s protocol, a pool of 3 plasmids, each encoding the Cas9 nuclease and target-specific 20 nt guide RNA were transfected into the cells. Successful transfection was confirmed by detection of GFP. Cells were examined for CD47 expression by FACS analysis and sorted to obtain cells showing no CD47 expression. CRISPR/Cas9 Plasmid (Santa Cruz Biotechnology; catalog #sc-418922) containing non-targeting 20 nt scramble guide RNA was used as a control. Cells expressing GFP were selected, expanded, and sorted to obtain cells lacking CD47. The lack of CD47 was confirmed by Western blotting. The expression of Mac-1 in transfected cells was controlled by FACS analysis using mAb 44a. The HEK293-based cell variants were maintained in DMEM/F12 medium (10-092-CM, Corning, Manassas, VA) supplemented with 10% fetal bovine serum (FBS) and antibiotics. IC-21 murine macrophages were grown in RPMI (Corning; catalog #10-040-CM) containing 10% FBS and antibiotics. Resident peritoneal macrophages were obtained from eight-week-old WT and CD47^-/-^ mice by lavage using cold PBS containing 5 mM EDTA as described (34). Inflammatory macrophages were collected 3 days after TG injection by peritoneal lavage with 5 mL ice-cold PBS+5 mM EDTA. Macrophages were isolated from the peritoneal lavage using the EasySep Mouse selection kit (StemCell Technologies, Vancouver, BC, Canada) with mAb F4/80 conjugated to PE. Human blood monocytes were isolated from the PBMC fraction using EasySEP Human Monocyte Isolation Kit.

### Expression of recombinant proteins

The cDNA encoding the extracellular N-terminal IgV domain of human CD47 (residues 19-141) was amplified from the plasmid hCD47 VersaClone cDNA (RDC1523, R&D systems) and was cloned in the expression vector pGEX-4T-1 (GE Healthcare). Recombinant CD47 was expressed as a fusion protein with glutathione *S*-transferase (GST) and purified from soluble fractions of *E. coli* lysates by affinity chromatography using Glutathione-agarose.

Recombinant Mac-1-derived fragments spanning the human extracellular domains EGF-3 (residues 513-551), EGF-4 (residues 552-596), βT (residues 597-673), EGF-3-4-βT (residues 513-673) and EGF-4-βT (residues 552-673) of the integrin β_2_ subunit; calf-1 (residues 764-901) and calf-2 (residues 902-1086) of the α_M_ subunit were expressed as fusion proteins with a polyhistidine tag. The coding regions were amplified by PCR using the full-length cDNA of α_M_ and β_2_ subunits as a template. The fragments were digested with appropriate restriction enzymes and cloned in the pET15b expression vector (EMD Millipore, Billerica, MA). The plasmids were transformed in *E. coli* strain BL-21(DE3)pLysS-competent cells, and expression was induced by adding 1 mM isopropyl 1-thio-β-d-galactopyranoside for 5 h at 37 °C. Proteins were purified using Ni-NTA from soluble fractions of *E. coli* lysates by metal-affinity chromatography. The fragments spanning the IgV domains of mouse SIRPa were previously described (22).

### Cell adhesion assay

Adhesion assays were performed as described previously (36,64). Briefly, the wells of 96-well microtiter plates (Immunol 4HBX, catalog #3855, Thermo Fisher) were coated with various concentrations of fibrinogen or ICAM-1 for 3 h at 37 °C and post-coated with 1.0% PVP for 1 h at 37 °C. Cells were labeled with 5 μM calcein for 30 min at 37 °C and washed twice with Hanks’ Balanced Salt Solution (HBSS) containing 0.1% BSA. Aliquots (100 μl) of labeled cells (5×10^5^/ml) were added to each well and allowed to adhere for 30 min at 37 °C. The nonadherent cells were removed by two washes with PBS. Fluorescence was measured in a CytoFluorII fluorescence plate reader. In inhibition adhesion experiments, cells were mixed with different concentrations of recombinant CD47-derived IgV fragment for 20 min at 22 °C before they were added to the wells coated with 2.5 μg/ml fibrinogen.

### Cell spreading assay

To determine cell spreading, inflammatory macrophages isolated from the peritoneum of WT and CD47^-/-^ mice were allowed to adhere to glass coverslips coated with fibrinogen (2.5 μg/ml) or ICAM-1 (2 μg/ml) for 1 h at 37 °C. Macrophages were fixed with 2% paraformaldehyde, permeabilized with 0.1% Triton X-100, and incubated with Alexa Fluor 488-conjugated phalloidin and DAPI. Confocal images were obtained with a Leica SP8 Confocal System (Exton, PA) using a 63x/1.4 objective. The cell area was calculated using ImageJ software (NIH, Bethesda, MD).

### Transwell migration assay

Migration assays with purified inflammatory macrophages isolated from the peritoneum of WT and CD47^-/-^ mice using Transwell inserts (5 μm pore size) were performed as previously described (32,33). Briefly, the lower chamber of the Transwell system contained 600 μl of 5 μg/ml endotoxin-free LL-37. F4/80-PE-labeled macrophages (100 μl at 3 × 10^6^ /ml) were added to the upper wells of the Transwell chamber and allowed to migrate for 90 min at 37 °C in a 5% CO_2_ humidified atmosphere. The assay was stopped by removing cells from the upper part of the filter separating the two chambers. Cells migrating to the bottom of the filter were detected using a Leica DM4000 B microscope (Leica Microsystems, Buffalo Grove, IL).

### Phagocytosis assay

Phagocytosis of fluorescent beads by adherent macrophages was previously described (32,33). Briefly, inflammatory macrophages isolated from the peritoneum of WT and CD47^-/-^ mice were allowed to adhere to glass coverslips for 2 h at 37 °C. After removing nonadherent cells, fluorescent latex beads coated with the cathelicidin peptide LL-37 were added to the cells and incubated for 30 min at 37 °C. Cells were washed with PBS, fixed with 2% paraformaldehyde, and beads were counted in the presence of trypan blue. The ratio of bacterial particles per macrophage was quantified by taking photographs of three fields for each well using a Leica DM4000 B microscope (Leica Microsystems, Buffalo Grove, IL) with a 20x objective.

### Macrophage fusion

Macrophage fusion was induced as previously described (66). Briefly, inflammatory macrophages isolated from the peritoneum of WT and CD47^-/-^ mice were allowed to adhere to paraffin-coated glass coverslips. After incubation in 5% CO_2_ at 37 °C for 30 min, nonadherent cells were removed, and adherent cells were continued to culture in complete DMEM/F12 media with 10% FBS for 2 h. Fusion was induced by adding 10 ng/ml IL-4 to the media. Cells were cultured for 6, 12, 24, and 48 h after IL-4 addition. After indicated time points, cells were fixed with 3.7% paraformaldehyde, followed by staining with Alexa Fluor 647-conjugated phalloidin and 4,6-diamidino-2-phenylindole (DAPI). Images were obtained with a Leica SP8 Confocal System (Exton, PA), and fusion index was calculated from images as described (66).

### Immunoprecipitation

Mac-1-HEK293 cells, HEK293 cells expressing the I-less form of Mac-1 or individual integrin subunits, IC-21 macrophages, inflammatory peritoneal macrophages, and human monocytes were solubilized with a lysis buffer (20 mM Tris-HCl, pH 7.4, 150 mM NaCl, 1% Triton X-100, 1 mM CaCl2, 1 mM PMSF, and protease inhibitor cocktail for 30 min at 22 °C. After removing insoluble material by centrifugation at 12000*g* for 15 min, the lysates were incubated with 10 *μ*g of normal mouse IgG and 50 *μ*L of Zysorbin-G for 2 h at 4 °C. After centrifugation, the supernatants were incubated with mAb 44A, anti-mouse α_M_ polyclonal antibody, mAb IB4, anti-β_1_ mAb 1965, anti-human/mouse β_1_ polyclonal antibody, mAb B6H12, or anti-human/mouse CD47 polyclonal antibody for 2 h at 4 °C. The integrin-mAb complexes were then collected by incubating with 50 *μ*L of protein A-Sepharose overnight at 4 °C. The immunoprecipitated proteins were eluted with SDS-PAGE loading buffer, electrophoresed on 7.5% SDS-polyacrylamide gels, and analyzed by Western blotting. To detect the presence of selected proteins, Immobilon P membranes were incubated with anti-human α_M_ mAb 44a (1:2000), polyclonal anti-mouse α_M_ (1 μg/ml), polyclonal anti-human/mouse β_1_ (1:2000), polyclonal anti-human/mouse CD47 (1:500), mAb B6H12 (5 μg/ml) and developed using SuperSignal West Pico substrate (ThermoFisher). In some experiments, cells (5×10^6^/0.5 ml) were labeled with 100 μg Immunopure Sulfo-NHS-LC-Biotin (Thermo Scientific, Rockford, IL) for 30 min at 22 °C. After solubilization and immunoprecipitation, the Immobilon P membranes were incubated with horseradish peroxidase-conjugated streptavidin and developed using SuperSignal West Pico substrate.

### Solid Phase Binding Assays

To test the interaction of IgV-CD47 with the β_2_ and α_M_-derived recombinant fragments, 96-well plates (Immulon 4BX, Thermo Scientific Rockford, IL) were coated with IgV-CD47 freed from its fusion part (2 μg/ml) overnight at 4 °C and postcoated with 1% BSA for 1 h at 22 °C. The His-tagged fragments in 20 mM Tris-HCl, pH 7.4, 100 mM NaCl, 1 mM MgCl_2_, 1 mM CaCl_2_, 0.05% Tween 20 were added to the wells and incubated for 3 h at 37 °C. After washing, bound proteins were detected using an anti-His mAb (5 μg/ml). After washing, goat anti-mouse IgG conjugated to alkaline phosphatase (AP) was added for 1 h, and the binding of recombinant proteins was measured by reaction with *p*-nitrophenyl phosphate. Background binding to BSA was subtracted.

### Flow Cytometry

FACS analyses were performed to assess the expression of α_M_ and β_2_ integrin subunits on the surface of transfected HEK293. Cells were incubated with anti-α_M_ mAb 44a (10 μg/ml) or anti-β_2_ mAb IB4 (10 μg/ml) followed by Alexa Fluor 488-conjugated secondary antibody and analyzed using a FACScan (BD Biosciences) and Attune NxT flow cytometer (ThermoFisher). To assess the expression of the epitope for mAb KIM127, Mac-1-HEK293 cells were activated with Mn^2+^ (1 mM) and PMA (0.1 μM) for 10 min at 22 °C and incubated with mAb KIM127 10 μg/ml for 30 min at 37 °C followed by Alexa Fluor 488-conjugated secondary antibody. The KIM127 epitope expression after incubation of Mac-1-HEK293 cells with mAb MEM48 was assayed using FITC-conjugated KIM127.

### Statistical analysis

All data are presented as the mean ± SE. The statistical differences between the two groups were determined using a Student’s t-test. Multiple comparisons were made using ANOVA followed by Tukey’s or Dunn’s post-test using GraphPad Instat software. Differences were considered significant at *P* < 0.05.

## Acknowledgment

This work was supported by the NIH grants HL63199 (TU) and GM118518 (XW) and the NSF GRFP grant 026257-001 (SK). We acknowledge the use of instruments within the Biosciences Advances Light Microscopy Facility at Arizona State University. Image data were collected using a Leica TCS SP5 LSCM (the National Institutes of Health SIG award S10 RR027154) and a Leica TCS SP8 LSCM (the NIH SIG award S10 OD023691).

## SUPPLEMENTAL FIGURE LEGENDS

**Supplemental Figure 1. Expression of Mac-1 and CD47 in WT and CD47-deficient macrophages.** Inflammatory macrophages were isolated from the peritoneum of WT and CD47-/- mice three days after TG injection and analyzed by FACS with a rat mAb against mouse CD47 (upper panels) and mAb M1/70 against the mouse α_M_ integrin subunit (bottom panels). Blue histograms indicate cells incubated with primary and secondary antibodies, and red histograms show cells incubated with secondary antibodies (control). MFI, mean fluorescence intensity. n=3.

**Supplemental Figure 2. Effect of activation on adhesion of Mac-1-HEK293 cells to fibrinogen.** Aliquots of calcein-labeled and activated by Mn^2+^ and PMA Mac-1-HEK293 cells were added to microtiter wells coated with 5 μg/ml of fibrinogen. After 30 min at 37 °C, nonadherent cells were removed by washing, and the fluorescence of adherent cells was measured. Results are normalized to the adhesion of nonactivated cells. Data are means ± SE from 3 experiments with triplicate measurements. ***p < .001.

**Supplemental Figure 3. Adhesion of WT and CD47-deficient Mac-1-HEK293 cells.** Aliquots of calcein-labeled WT and CD47 KO macrophages were activated by Mn^2+^ and added to microtiter wells coated with 5 μg/ml of fibrinogen. After 30 min at 37 °C, nonadherent cells were removed by washing, and the fluorescence of adherent cells was measured. Data are expressed as a percentage of adhesion and are means ± SE from 3 experiments with triplicate measurements. **p≤ 0.01.

## REFERENCES

1. Brown, E. J., and Frazier, W. A. (2001) Integrin-associated protein (CD47) and its ligands. Trends Cell Biol. 11, 130–135

2. Oldenborg, P. A. (2013) CD47: A Cell Surface Glycoprotein Which Regulates Multiple Functions of Hematopoietic Cells in Health and Disease. ISRN.Hematol. 2013, 614–619

3. Soto-Pantoja, D. R., Kaur, S., and Roberts, D. D. (2015) CD47 signaling pathways controlling cellular differentiation and responses to stress. Crit. Rev. Biochem. Mol. Biol. 50, 212–230

4. Sick, E., Jeanne, A., Schneider, C., Dedieu, S., Takeda, K., and Martiny, L. (2012) CD47 update: a multifaceted actor in the tumour microenvironment of potential therapeutic interest. Br. J. Pharmacol. 167, 1415–1430

5. Gresham, H. D., Goodwin, J. L., Allen, P. M., Anderson, D. C., and Brown, E. J. (1989) A novel member of the integrin receptor family mediates Arg-Gly-Asp-stimulated neutrophil phagocytosis. J. Cell Biol. 108, 1935–1943

6. Brown, E., Hooper, L., Ho, T., and Gresham, H. (1990) Integrin-associated protein: a 50-kD plasma membrane antigen physically and functionally associated with integrins. J. Cell Biol. 111, 2785–2794

7. Lindberg, F. P., Gresham, H. D., Schwarz, E., and Brown, E. J. (1993) Molecular cloning of integrin-associated protein: an immunoglobulin family member with multiple membrane-spanning domains implicated in alpha v beta 3-dependent ligand binding. J. Cell Biol. 123, 485–496

8. Senior, R. M., Gresham, H. D., Griffin, G. L., Brown, E. J., and Chung, A. E. (1992) Entactin stimulates neutrophil adhesion and chemotaxis through interactions between its Arg-Gly-Asp (RGD) domain and the leukocyte response integrin. J. Clin. Invest. 90, 2251–2257

9. Schwartz, M. A., Brown, E. J., and Fazeli, B. (1993) A 50-kDa integrin-associated protein is required for integrin-regulated calcium entry in endothelial cells. J. Biol. Chem. 268, 19931–19934

10. Parkos, C. A., Colgan, S. P., Liang, T. W., Nusrat, A., Bacarra, A. E., Carnes, D. K., and Madara, J. L. (1996) CD47 mediates post-adhesive events required for neutrophil migration across polarized intestinal epithelia. J. Cell Biol. 132, 437–450

11. Cooper, D., Lindberg, F. P., Gamble, J. R., Brown, E. J., and Vadas, M. A. (1995) Transendothelial migration of neutrophils involves integrin-associated protein (CD47). Proc. Natl. Acad. Sci. U. S. A. 92, 3978–3982

12. Lindberg, F. P., Gresham, H. D., Reinhold, M. I., and Brown, E. J. (1996) Integrin-associated protein immunoglobulin domain is necessary for efficient vitronectin bead binding. J. Cell Biol. 1313, 1322

13. Lindberg, F. P., Bullard, D. C., Caver, T. E., Gresham, H. D., Beaudet, A. L., and Brown, E. J. (1996) Decreased resistance to bacterial infection and granulocyte defects in IAP-deficient mice. Science 274, 795–798

14. Azcutia, V., Stefanidakis, M., Tsuboi, N., Mayadas, T., Croce, K. J., Fukuda, D., Aikawa, M., Newton, G., and Luscinskas, F. W. (2012) Endothelial CD47 promotes vascular endothelial-cadherin tyrosine phosphorylation and participates in T cell recruitment at sites of inflammation in vivo. J. Immunol. 189, 2553–2562

15. Chung, J., Wang, X. Q., Lindberg, F. P., and Frazier, W. A. (1999) Thrombospondin-1 acts via IAP/CD47 to synergize with collagen in alpha2beta1-mediated platelet activation. Blood 94, 642–648

16. Fujimoto, T. T., Katsutani, S., Shimomura, T., and Fujimura, K. (2003) Thrombospondin-bound integrin-associated protein (CD47) physically and functionally modifies integrin alphaIIbbeta3 by its extracellular domain. J. Biol. Chem. 278, 26655–26665

17. Wang, X. Q., and Frazier, W. A. (1998) The thrombospondin receptor CD47 (IAP) modulates and associates with alpha2 beta1 integrin in vascular smooth muscle cells. Mol. Biol. Cell 9, 865–874

18. Brittain, J. E., Han, J., Ataga, K. I., Orringer, E. P., and Parise, L. V. (2004) Mechanism of CD47-induced alpha4beta1 integrin activation and adhesion in sickle reticulocytes. J. Biol. Chem. 279, 42393–42402

19. Orazizadeh, M., Lee, H. S., Groenendijk, B., Sadler, S. J., Wright, M. O., Lindberg, F. P., and Salter, D. M. (2008) CD47 associates with alpha 5 integrin and regulates responses of human articular chondrocytes to mechanical stimulation in an in vitro model. Arthritis Res. Ther. 10, R4

20. Azcutia, V., Routledge, M., Williams, M. R., Newton, G., Frazier, W. A., Manica, A., Croce, K. J., Parkos, C. A., Schmider, A. B., Turman, M. V., Soberman, R. J., and Luscinskas, F. W. (2013) CD47 plays a critical role in T-cell recruitment by regulation of LFA-1 and VLA-4 integrin adhesive functions. Mol. Biol. Cell 24, 3358–3368

21. Ticchioni, M., Raimondi, V., Lamy, L., Wijdenes, J., Lindberg, F. P., Brown, E. J., and Bernard, A. (2001) Integrin-associated protein (CD47/IAP) contributes to T cell arrest on inflammatory vascular endothelium under flow. FASEB J. 15, 341–350

22. Podolnikova, N. P., Hlavackova, M., Yakubenko, V. P., Faust, J. J., Balabiyev, A., Wang, X., and Ugarova, T. P. (2019) Interaction between the integrin Mac-1 and SIRPa mediates fusion in heterologous cells. J. Biol. Chem. 294(19), 7833–7849

23. Coxon, A., Rieu, P., Barkalow, F. J., Askari, S., Sharpe, A. H., Von Andrian, U. H., Arnaout, M. A., and Mayadas, T. N. (1996) A novel role for the beta 2 integrin CD11b/CD18 in neutrophil apoptosis: a homeostatic mechanism in inflammation. Immunity 5, 653–666

24. Lu, H., Smith, C. W., Perrard, J., Bullard, D., Tang, L., Entman, M. L., Beaudet, A. L., and Ballantyne, C. M. (1997) LFA-1 is sufficient in mediating neutrophil emigration in Mac-1 deficient mice. JCI 99, 1340–1350

25. Campbell, I. D., and Humphries, M. J. (2011) Integrin structure, activation, and interactions. Cold Spring Harb. Perspect. Biol. 3

26. Luo, B. H., Carman, C. V., and Springer, T. A. (2007) Structural basis of integrin regulation and signaling. Annu. Rev. Immunol. 25, 619–647

27. Flick, M. J., Du, X., Witte, D. P., Jirousková, M., Soloviev, D. A., Busuttil, S. J., Plow, E. F., and Degen, J. L. (2004) Leukocyte engagement of fibrin(ogen) via the integrin receptor alphaMbeta2/Mac-1 is critical for host inflammatory response in vivo. J. Clin. Invest. 113, 1596–1606

28. Diamond, M. S., Staunton, D. E., de Fougerolles, A. R., Stacker, S. A., Garcia-Aguilar, J., Hibbs, M. L., and Springer, T. A. (1990) ICAM-1 (CD54)-a counter-receptor for MAC-1 (CD11b/CD18). J. Cell Biol. 111, 3129–3139

29. Lishko, V. K., Burke, T., and Ugarova, T. P. (2007) Anti-adhesive effect of fibrinogen: A safeguard for thrombus stability. Blood 109, 1541–1549

30. Podolnikova, N. P., Yermolenko, I. S., Fuhrmann, A., Lishko, V. K., Magonov, S., Bowen, B., Enderlein, J., Podolnikov, A., Ros, R., and Ugarova, T. P. (2010) Control of integrin aIIbb3 outside-in signaling and platelet adhesion by sensing the physical properties of fibrin(ogen) substrates Biochemistry 49, 68–77

31. Cao, C., Lawrence, D. A., Strickland, D. K., and Zhang, L. (2005) A specific role of integrin Mac-1 in accelerated efflux to the lymphatics. Blood 106, 3234–3241

32. Lishko, V. K., Moreno, B., Podolnikova, N. P., and Ugarova, T. P. (2016) Identification of human cathelicidin peptide LL-37 as a ligand for macrophage integrin α_M_β_2_ (Mac-1, CD11b/CD18) that promotes phagocytosis by opsonizing bacteria. Res. Rep. Biochem. 2016, 39–55

33. Lishko, V. K., Yakubenko, V. P., Ugarova, T. P., and Podolnikova, N. P. (2018) Leukocyte integrin Mac-1 (CD11b/CD18, α_M_β_2_, CR3) acts as a functional receptor for platelet factor 4. J. Biol. Chem. 293, 6869–6882

34. Podolnikova, N. P., Kushchayeva, Y. S., Wu, Y., Faust, J., and Ugarova, T. P. (2016) The Role of Integrins α_M_β_2_ (Mac-1, CD11b/CD18) and α_D_β_2_ (CD11d/CD18) in Macrophage Fusion. Am. J. Pathol. 186, 2105–2116

35. Faust, J. J., Christenson, W., Doudrick, K., Heddleston, J., Chew, T. L., Lampe, M., Balabiyev, A., Ros, R., and Ugarova, T. P. (2018) Fabricating Optical-quality Glass Surfaces to Study Macrophage Fusion. J.Vis.Exp. 133

36. Lishko, V. K., Yakubenko, V. P., and Ugarova, T. P. (2003) The interplay between Integrins α_M_β_2_ and α_5_β_1_ during cell migration to fibronectin. Exp. Cell Res. 283, 116–126

37. Lu, C., Ferzly, M., Takagi, J., and Springer, T. A. (2001) Epitope mapping of antibodies to the C-terminal region of the integrin beta 2 subunit reveals regions that become exposed upon receptor activation. J. Immunol. 166, 5629–5637

38. Nishida, N., Walz, T., and Springer, T. A. (2006) Structural transitions of complement component C3 and its activation products. Proc. Natl. Acad. Sci. U. S. A. 103, 19737–19742

39. Xie, C., Zhu, J., Chen, X., Mi, L., Nishida, N., and Springer, T. A. (2010) Structure of an integrin with an alphaI domain, complement receptor type 4. EMBO J. 29, 666–679

40. Fenalti, G., Villanueva, N., Griffith, M., Pagarigan, B., Lakkaraju, S. K., Huang, R. Y., Ladygina, N., Sharma, A., Mikolon, D., Abbasian, M., Johnson, J., Hadjivassiliou, H., Zhu, D., Chamberlain, P. P., Cho, H., and Hariharan, K. (2021) Structure of the human marker of self 5-transmembrane receptor CD47. Nat Commun 12, 5218

41. Rebres, R. A., Vaz, L. E., Green, J. M., and Brown, E. J. (2001) Normal ligand binding and signaling by CD47 (integrin-associated protein) requires a long range disulfide bond between the extracellular and membrane-spanning domains. J. Biol. Chem. 276, 34607–34616

42. Xiong, J. P., Stehle, T., Diefenbach, B., Zhang, R., Dunker, R., Scott, D. L., Joachimiak, A., Goodman, S. L., and Arnaout, M. A. (2001) Crystal structure of the extracellular segment of integrin alpha v beta 3 Science 294, 339–345

43. Takagi, J., Petre, B., Walz, T., and Springer, T. A. (2002) Global conformational rearrangements in integrin extracellular domains in outside-in and inside-out signaling. Cell 110, 599–611

44. Springer, T. A., and Dustin, M. L. (2012) Integrin inside-out signaling and the immunological synapse. Curr. Opin. Cell Biol. 24, 107–115

45. Kim, M., Carman, C. V., and Springer, T. A. (2003) Bidirectional transmembrane signaling by cytoplasmic domain separation in integrins. Science 301, 1720–1725

46. Tang, R. H., Tng, E., Law, S. K., and Tan, S. M. (2005) Epitope mapping of monoclonal antibody to integrin alphaL beta2 hybrid domain suggests different requirements of affinity states for intercellular adhesion molecules (ICAM)-1 and ICAM-3 binding. J. Biol. Chem. 280, 29208–29216

47. Su, Y., Xia, W., Li, J., Walz, T., Humphries, M. J., Vestweber, D., Cabañas, C., Lu, C., and Springer, T. A. (2016) Relating conformation to function in integrin α5β1. Proc. Natl. Acad. Sci. U. S. A. 113, E3872–3881

48. Jiang, P., Lagenaur, C. F., and Narayanan, V. (1999) Integrin-associated protein is a ligand for the P84 neural adhesion molecule. J. Biol. Chem. 274, 559–562

49. Seiffert, M., Cant, C., Chen, Z., Rappold, I., Brugger, W., Kanz, L., Brown, E. J., Ullrich, A., and Bühring, H. J. (1999) Human signal-regulatory protein is expressed on normal, but not on subsets of leukemic myeloid cells and mediates cellular adhesion involving its counterreceptor CD47. Blood 94, 3633–3643

50. Oldenborg, P. A., Zheleznyak, A., Fang, Y. F., Lagenaur, C. F., Gresham, H. D., and Lindberg, F. P. (2000) Role of CD47 as a marker of self on red blood cells. Science 288, 2051–2054

51. Matozaki, T., Murata, Y., Okazawa, H., and Ohnishi, H. (2009) Functions and molecular mechanisms of the CD47-SIRPalpha signalling pathway. Trends Cell Biol. 19, 72–80

52. Bian, Z., Shi, L., Guo, Y. L., Lv, Z., Tang, C., Niu, S., Tremblay, A., Venkataramani, M., Culpepper, C., Li, L., Zhou, Z., Mansour, A., Zhang, Y., Gewirtz, A., Kidder, K., Zen, K., and Liu, Y. (2016) Cd47-Sirpα interaction and IL-10 constrain inflammation-induced macrophage phagocytosis of healthy self-cells. Proc. Natl. Acad. Sci. U. S. A. 113, E5434–5443

53. Logtenberg, M. E. W., Scheeren, F. A., and Schumacher, T. N. (2020) The CD47-SIRPα Immune Checkpoint. Immunity 52, 742–752

54. Jaiswal, S., Jamieson, C. H., Pang, W. W., Park, C. Y., Chao, M. P., Majeti, R., Traver, D., van Rooijen, N., and Weissman, I. L. (2009) CD47 is upregulated on circulating hematopoietic stem cells and leukemia cells to avoid phagocytosis. Cell 138, 271–285

55. Majeti, R., Chao, M. P., Alizadeh, A. A., Pang, W. W., Jaiswal, S., Gibbs, K. D., Jr., van Rooijen, N., and Weissman, I. L. (2009) CD47 is an adverse prognostic factor and therapeutic antibody target on human acute myeloid leukemia stem cells. Cell 138, 286–299

56. Willingham, S. B., Volkmer, J. P., Gentles, A. J., Sahoo, D., Dalerba, P., Mitra, S. S., Wang, J., Contreras-Trujillo, H., Martin, R., Cohen, J. D., Lovelace, P., Scheeren, F. A., Chao, M. P., Weiskopf, K., Tang, C., Volkmer, A. K., Naik, T. J., Storm, T. A., Mosley, A. R., Edris, B., Schmid, S. M., Sun, C. K., Chua, M. S., Murillo, O., Rajendran, P., Cha, A. C., Chin, R. K., Kim, D., Adorno, M., Raveh, T., Tseng, D., Jaiswal, S., Enger, P., Steinberg, G. K., Li, G., So, S. K., Majeti, R., Harsh, G. R., van de Rijn, M., Teng, N. N., Sunwoo, J. B., Alizadeh, A. A., Clarke, M. F., and Weissman, I. L. (2012) The CD47-signal regulatory protein alpha (SIRPa) interaction is a therapeutic target for human solid tumors. Proc. Natl. Acad. Sci. U. S. A. 109, 6662–6667

57. Huang, Y., Ma, Y., Gao, P., and Yao, Z. (2017) Targeting CD47: the achievements and concerns of current studies on cancer immunotherapy. J. Thorac. Dis. 9, E168–e174

58. Liu, X., Kwon, H., Li, Z., and Fu, Y. X. (2017) Is CD47 an innate immune checkpoint for tumor evasion? J. Hematol. Oncol. 10, 12

59. Oldenborg, P. A., Gresham, H. D., and Lindberg, F. P. (2001) CD47-signal regulatory protein alpha (SIRPalpha) regulates Fcgamma and complement receptor-mediated phagocytosis. J. Exp. Med. 193, 855–862

60. Aderem, A., and Underhill, D. M. (1999) Mechanisms of phagocytosis in macrophages. Annu. Rev. Immunol. 17, 593–623

61. Zhang, W., Huang, Q., Xiao, W., Zhao, Y., Pi, J., Xu, H., Zhao, H., Xu, J., Evans, C. E., and Jin, H. (2020) Advances in Anti-Tumor Treatments Targeting the CD47/SIRPα Axis. Front. Immunol. 11, 18

62. Blystone, S. D., Lindberg, F. P., LaFlamme, S. E., and Brown, E. J. (1995) Integrin beta 3 cytoplasmic tail is necessary and sufficient for regulation of alpha 5 beta 1 phagocytosis by alpha v beta 3 and integrin-associated protein. J. Cell Biol. 130, 745–754

63. Pietsch, E. C., Dong, J., Cardoso, R., Zhang, X., Chin, D., Hawkins, R., Dinh, T., Zhou, M., Strake, B., Feng, P. H., Rocca, M., Santos, C. D., Shan, X., Danet-Desnoyers, G., Shi, F., Kaiser, E., Millar, H. J., Fenton, S., Swanson, R., Nemeth, J. A., and Attar, R. M. (2017) Anti-leukemic activity and tolerability of anti-human CD47 monoclonal antibodies. Blood Cancer J 7, e536

64. Yakubenko, V. P., Lishko, V. K., Lam, S. C. T., and Ugarova, T. P. (2002) A molecular basis for integrin α_M_β_2_ ligand binding promiscuity J. Biol. Chem. 277, 48635–48642

65. Yalamanchili, P., Lu, C. F., Oxvig, C., and Springer, T. A. (2000) Folding and function of I domain-deleted Mac-1 and lymphocyte function-associated antigen-1. J. Biol. Chem. 275, 21877–21882

66. Faust, J. J., Balabiyev, A., Heddleston, J. M., Podolnikova, N. P., Baluch, D. P., Chew, T. L., and Ugarova, T. P. (2019) An actin-based protrusion originating from a podosome-enriched region initiates macrophage fusion. Mol. Biol. Cell 30, 2254–2267

