## Supplementary figures and images for "THE ASSOCIATION OF CD47 WITH INTEGRIN Mac-1 REGULATES MACROPHAGE RESPONSES BY STABILIZING THE EXTENDED INTEGRIN CONFORMATION"

### Supplemental Figure 1

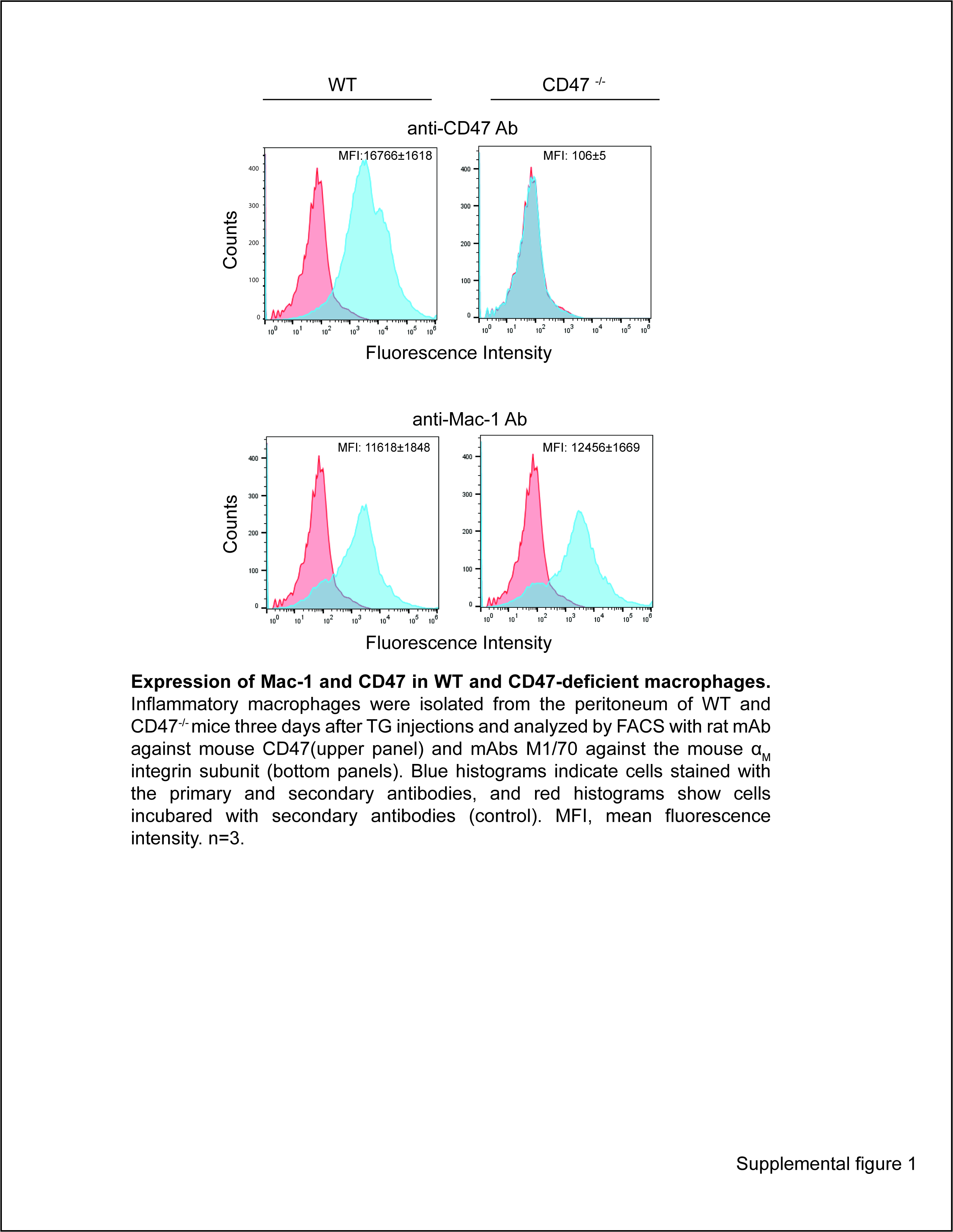

### Supplemental Figure 2

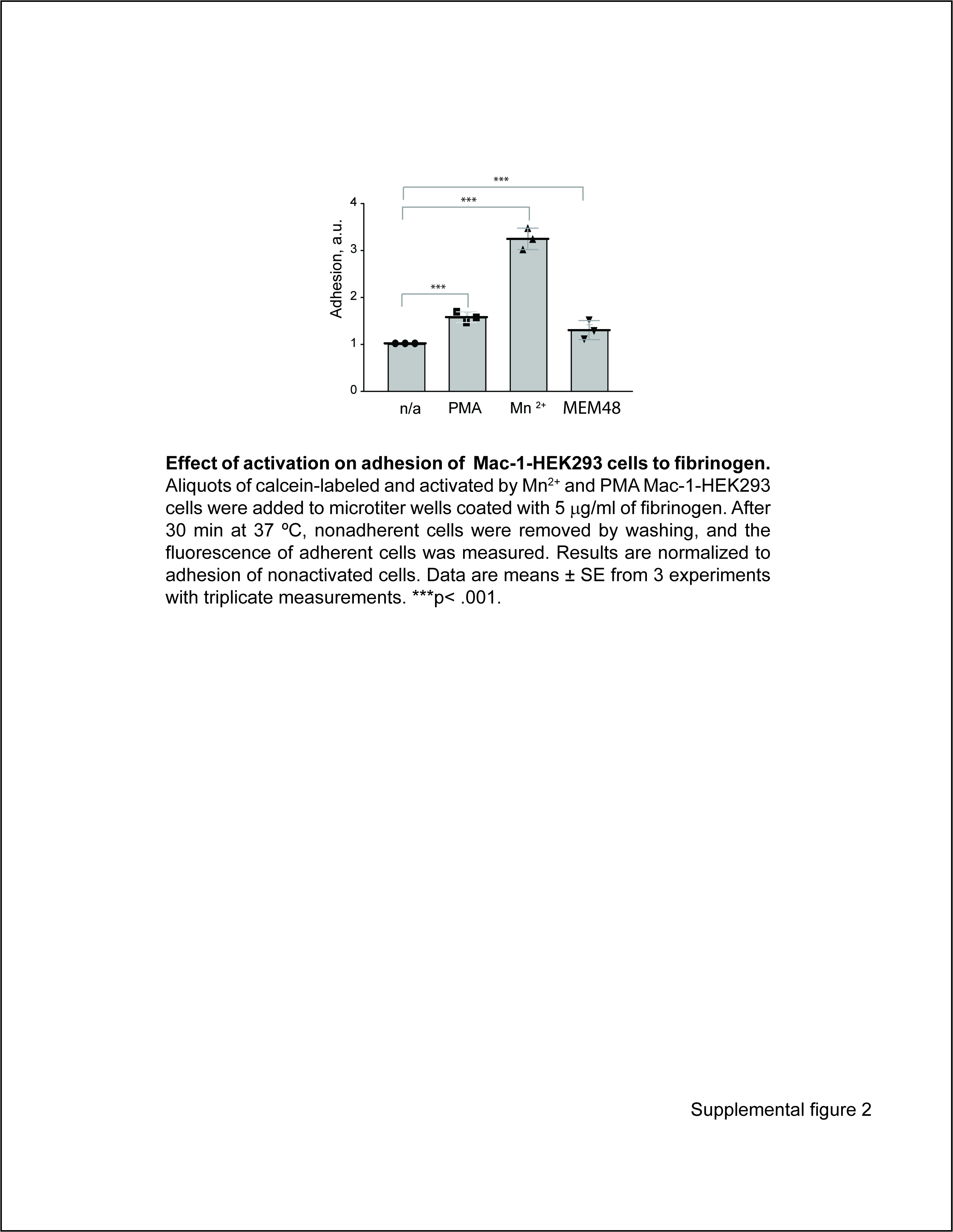

### Supplemental Figure 3

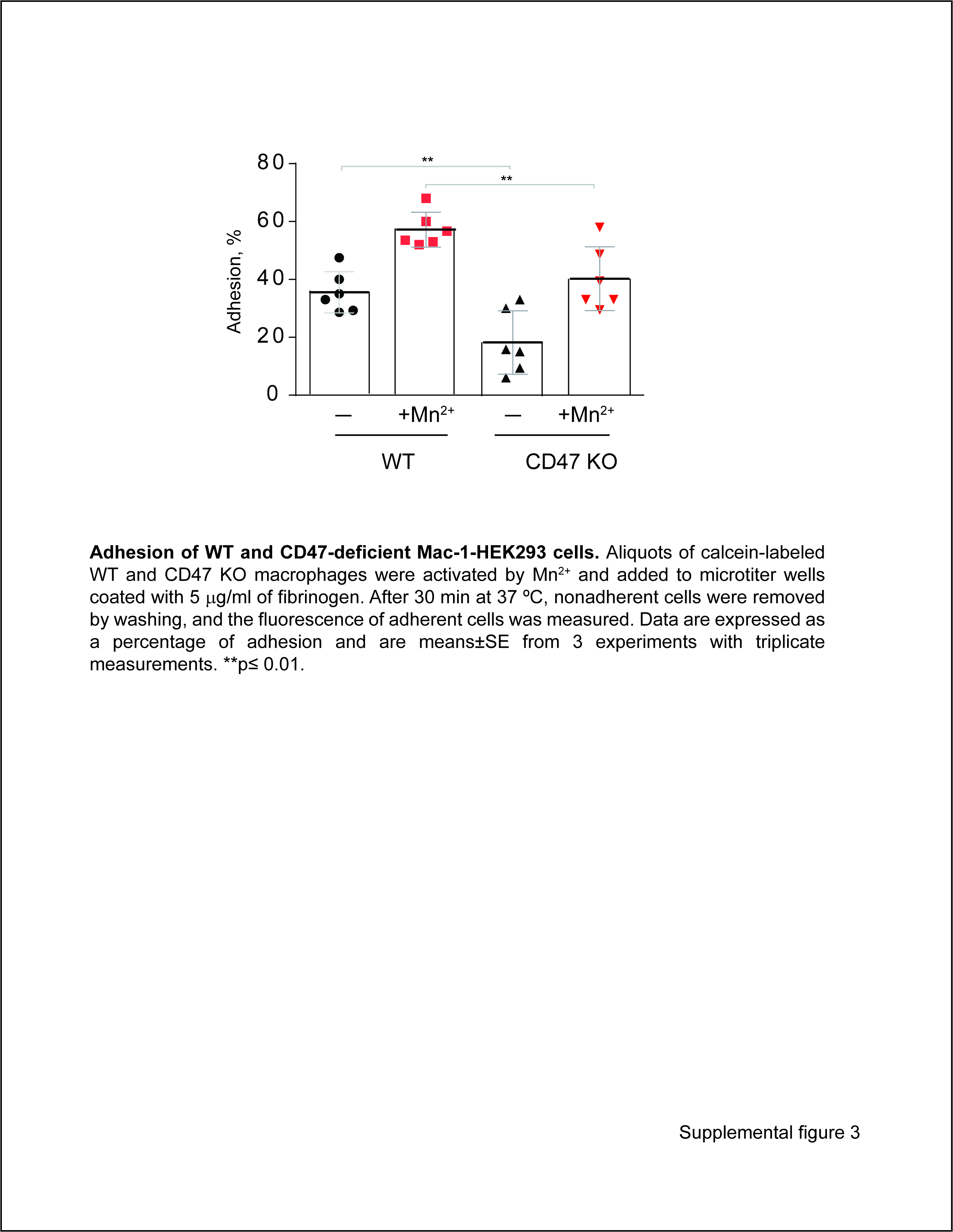
